# Targeting PIKfyve-driven lipid homeostasis as a metabolic vulnerability in pancreatic cancer

**DOI:** 10.1101/2024.03.18.585580

**Authors:** Caleb Cheng, Jing Hu, Rahul Mannan, Rupam Bhattacharyya, Nicholas J. Rossiter, Brian Magnuson, Jasmine P. Wisniewski, Yang Zheng, Lanbo Xiao, Chungen Li, Dominik Awad, Tongchen He, Yi Bao, Yuping Zhang, Xuhong Cao, Zhen Wang, Rohit Mehra, Pietro Morlacchi, Vaibhav Sahai, Marina Pasca di Magliano, Yatrik M. Shah, Ke Ding, Yuanyuan Qiao, Costas A. Lyssiotis, Arul M. Chinnaiyan

**Author notes:** Correspondence to: Arul M. Chinnaiyan, M.D., Ph.D., Costas A. Lyssiotis, Ph.D., and Yuanyuan Qiao, Ph.D.

## Abstract

Pancreatic ductal adenocarcinoma (PDAC) subsists in a nutrient-deregulated microenvironment, making it particularly susceptible to treatments that interfere with cancer metabolism^1^ ^2^. For example, PDAC utilizes and is dependent on high levels of autophagy and other lysosomal processes^3-5^. Although targeting these pathways has shown potential in preclinical studies, progress has been hampered by the challenge of identifying and characterizing favorable targets for drug development^6^. Here, we characterize PIKfyve, a lipid kinase integral to lysosomal functioning^7^, as a novel and targetable vulnerability in PDAC. In human patient and murine PDAC samples, we discovered that *PIKFYVE* is overexpressed in PDAC cells compared to adjacent normal cells. Employing a genetically engineered mouse model, we established the essential role of PIKfyve in PDAC progression. Further, through comprehensive metabolic analyses, we found that PIKfyve inhibition obligated PDAC to upregulate *de novo* lipid synthesis, a relationship previously undescribed. PIKfyve inhibition triggered a distinct lipogenic gene expression and metabolic program, creating a dependency on *de novo* lipid metabolism pathways, by upregulating genes such as *FASN* and *ACACA*. In PDAC, the KRAS-MAPK signaling pathway is a primary driver of *de novo* lipid synthesis, specifically enhancing *FASN* and *ACACA* levels. Accordingly, the simultaneous targeting of PIKfyve and KRAS-MAPK resulted in the elimination of tumor burden in a syngeneic orthotopic model and tumor regression in a xenograft model of PDAC. Taken together, these studies suggest that disrupting lipid metabolism through PIKfyve inhibition induces synthetic lethality in conjunction with KRAS-MAPK-directed therapies for PDAC.

## Main Text

Pancreatic ductal adenocarcinoma (PDAC) is one of the deadliest cancers with a five-year survival rate of just 13%^8^. This is mediated in large part by a lack of effective therapeutic options. The PDAC tumor microenvironment is central to this resistance and features a high degree of stromal fibroblasts and extracellular matrix deposition that cause PDAC to experience elevated interstitial pressures, low vascularity, and a disrupted nutritional availability^9^. To circumvent deregulated nutrient access, PDAC cells become expert scavengers, employing intra- and extracellular recycling pathways, sourcing non-classical nutrients from their environment through expression of high avidity nutrient transporters, bulk engulfment, and crosstalk with other pro-tumor cell types^3,5,9-15^.

Incidentally, these unique metabolic dependencies also provide opportunities for therapeutic interventions^1,16-18^. Specifically, previous studies support targeting lysosome-dependent pathways as a therapeutic strategy for PDAC^19^. Lysosome-dependent pathways serve multiple roles in PDAC^20^. For example, these pathways have been shown to maintain the availability of biosynthetic intermediates^33,5,11,21,22^, iron homeostasis^23-25^, and also to degrade MHC-1, increasing immune evasion^26,27^. These studies provided support for targeting autophagy and lysosome-dependent pathways to disrupt PDAC metabolism as a therapeutic strategy and have resulted in many clinical trials utilizing autophagy and lysosomal inhibitor hydroxychloroquine (HCQ) with chemotherapy in PDAC (NCT01273805, NCT01978184, NCT01506973, NCT04911816, NCT04524702, NCT01494155, NCT03344172)^28-30^.

Further highlighting this strategy were the findings that autophagy enables PDAC to adapt to inhibition of Kirsten rat sarcoma virus (KRAS) or the downstream mitogen-activated protein kinase (MAPK) pathway^31-34^. In nearly all cases of PDAC, KRAS harbors an activating mutation and drives metabolic homeostasis by signaling through the MAPK pathway^35^. While KRAS was, until recently, thought to be undruggable^36^, large-scale efforts to target KRAS^37^ have resulted in multiple direct inhibitors of KRAS that add to the existing arsenal of compounds targeting the MAPK pathway, especially MEK and ERK ^38-43^. In response to the findings that PDAC utilizes autophagy to adapt to KRAS-MAPK inhibition, two clinical trials are underway to actively investigate the safety and efficacy of combining MEK or ERK inhibitors with HCQ (NCT04386057, NCT04132505)^44,45^.

Despite the considerable interest and promise to target autophagy and lysosomal processes in PDAC, preclinical and clinical studies have been hampered by the lack of effective therapeutics targeting specific effectors of these processes^46^. The lipid kinase PIKfyve serves as the only cellular source of PI(3,5)P_2_ and PI5P, crucial phospholipids for lysosome activity^7^. Previous work illustrated that inhibition of PIKfyve disrupted autophagy flux and lysosome function, leading to increased immune activity and tumor suppression in multiple cancer models^47-51^. Importantly two PIKfyve inhibitors, apilimod and ESK981, have cleared phase 1 clinical trials (NCT02594384, NCT00875264)^52,53^, highlighting the rapid translational potential of targeting PIKfyve as a means to disrupt autophagy and lysosomal processes in cancers.

To this end, we sought to comprehensively characterize PIKfyve as a therapeutic target in PDAC. First, we identified that human and murine PDAC cells express more *PIKFYVE* transcripts than the surrounding normal pancreatic cells, a finding that has never been reported to our knowledge. We next utilized multiple approaches to perturb PIKfyve in PDAC models. Specifically, we generated a genetically engineered mouse model (GEMM) harboring a conditional deletion of *Pikfyve* and found that *Pikfyve* loss dramatically increased animal survival and decreased PDAC disease burden. Similarly, prophylactic pharmacological inhibition of PIKfyve also decreased PDAC disease burden in a GEMM of PDAC. To assess the metabolic impact of PIKfyve inhibition in PDAC, we performed a metabolism-focused CRISPR screen on PDAC cells. We discovered a synthetic dependency on *de novo* fatty acid synthesis through genes such as Fatty acid synthase (*FASN*) and Acetyl-CoA Carboxylase alpha (*ACACA*, protein name ACC1) following PIKfyve inhibition^54^. This relationship has not previously been described using other autophagy or lysosomal inhibitors^23,55^. Corroborating this observation using a multi-omics approach, we established that PIKfyve inhibition drives PDAC cells into a lipogenic transcriptional and metabolic state, suggesting lipid synthesis is a necessary adaptive process in response to PIKfyve inhibition.

In PDAC, FASN is overexpressed and has been nominated as a therapeutic target^56,57,58^. The revelation that PIKfyve inhibition uncovers *FASN* and *ACACA* as synthetic lethalities highlights PIKfyve as a promising therapeutic partner to inhibitors of the fatty acid synthesis pathway. In our study, we further find that KRAS-MAPK inhibition decreased expression of FASN and ACC1, establishing a novel relationship of synthetic dependency between PIKfyve and KRAS-MAPK in the regulation of lipid metabolism. Taken in the context of the previously established concept of autophagy as an adaptive mechanism for PDAC in response to KRAS-MAPK inhibition, our finding provides an additional, mechanistically distinct rationale for combining PIKfyve and KRAS-MAPK inhibitors. Thus, we tested this combinatorial regimen on multiple murine models and found that the dual inhibition resulted in sustained tumor regression or elimination, while each individual treatment had more modest effects. Taken together, our studies establish PIKfyve as a targetable metabolic vulnerability in PDAC and demonstrate that dual inhibition of PIKfyve and KRAS-MAPK is a promising and rapidly translatable therapeutic strategy for PDAC.

## Results

### *Pikfve* is dispensable for normal pancreas but is required for PDAC development

To study the role of *Pikfyve* in pancreatic cancer development, we first evaluated *Pikfyve* expression in the autochthonous PDAC GEMM *Pft1a-Cre; LSL-Kras^G12D/+^*; LSL-*Trp53^R172H/+^* (KPC). Employing BaseScope, an RNA *in situ* hybridization (RNA-ISH) technique with a short probe specifically targeting *Pikfyve* exon 6, we discovered that *Pikfyve* expression was dramatically and consistently higher in PanIN and PDAC tissue compared to surrounding normal tissue *in situ* **(Fig. 1A-B)**. We next assessed whether this overexpression was also seen in a panel of human PDAC samples archived at the University of Michigan Department of Pathology. Using RNA-ISH, we found that *PIKFYVE* was overexpressed in PDAC cells compared to matched, surrounding normal pancreatic cells (**Fig. 1C-D, Extended Data Fig. 1A**). These data suggested that PanIN and PDAC may have an elevated utilization of PIKfyve-driven processes, relative to normal pancreatic tissue. To assess this, we first generated conditional pancreatic *Pikfyve* knockout mice using the *Ptf1a* promoter-driven Cre recombinase (*Ptf1a-Cre; Pikfyve^f/f^)* **(Fig. 1E)**. Upon confirming the loss of PIKfyve protein in pancreatic tissue **(Fig. 1F-G)**, we assessed the physiological impact of *Pikfyve* loss on pancreatic development. We found that *Pikfyve* loss did not impact pancreatic weight, morphology, or function in terms of insulin production **(Extended Data Fig. 1B-C)**, suggesting that *Pikfyve* is not critical for normal pancreatic tissue development or function.

**Figure 1:**
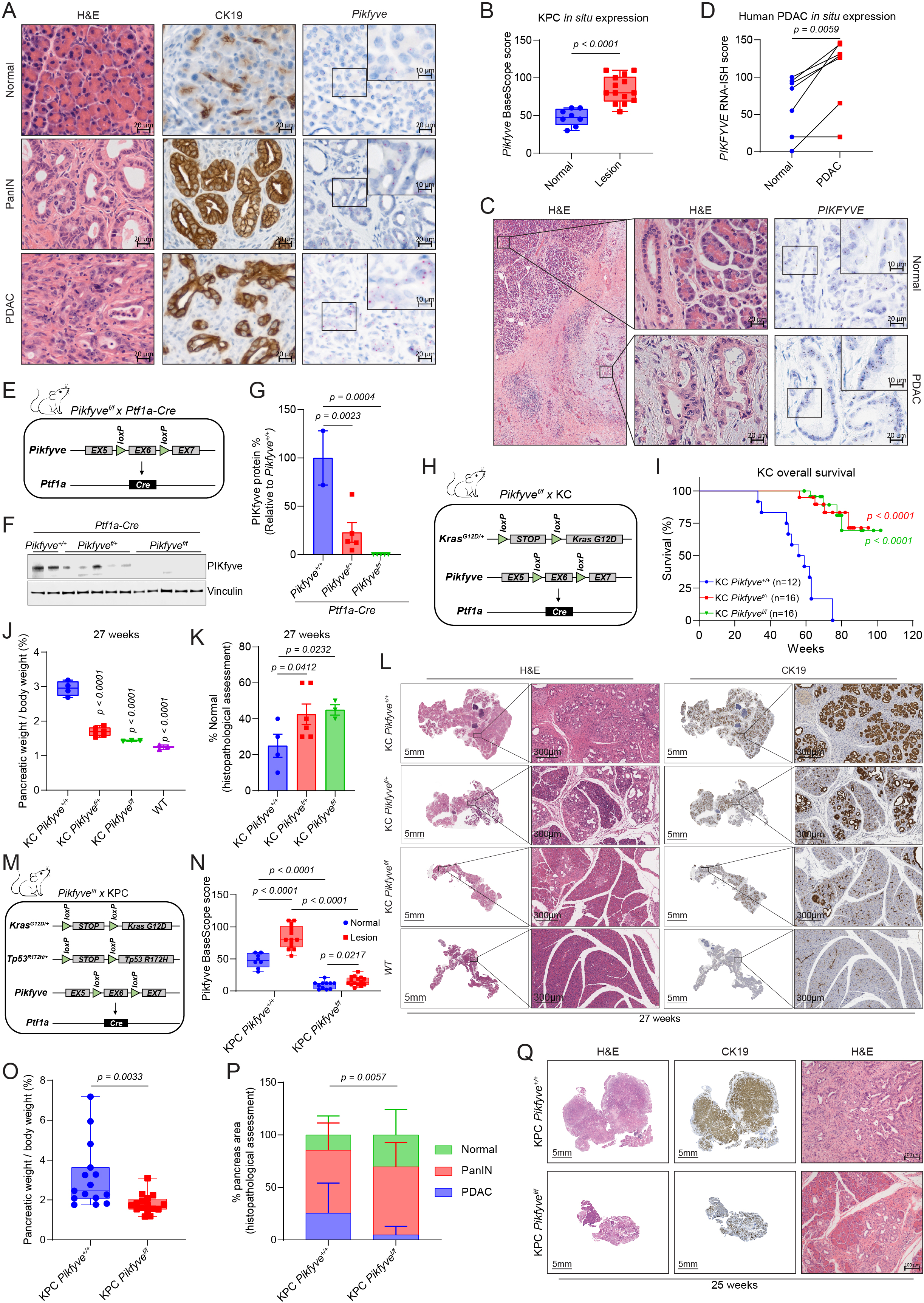
*Pikfyve* is essential for progression of precursor PanIN lesions to PDAC. A. Representative images of PanIN or PDAC lesions and normal tissue taken from a KPC murine pancreas, showing H&E, IHC staining for CK19, and BaseScope for *Pikfyve*. Scalebar = 20μm for low-magnification images and 10μm for high-magnification images. B. *In situ Pikfyve* levels in KPC murine pancreas lesion (PanIN or PDAC) vs normal tissue as determined by BaseScope RNA-ISH probes targeting *Pikfyve* exon 6. (Unpaired two-tailed t-test) C. Representative image of one human PDAC patient sample, showing H&E (left and middle sections) or *PIKFYVE* RNA-ISH (right). Scalebars are 200μm (left), 20μm (middle), 20μm (right, low magnification), and 10μm (right, high magnification). D. *In situ PIKFYVE* levels in histologically normal or PDAC cells in seven human PDAC patient samples using RNA-ISH (RNAScope). Biospy samples were taken from five independent PDAC patients: two patients donated two samples each from distinct biopsies. Scores were determined as described in methods. (Paired two-tailed t-test) E. Breeding design for the generation of *Pikfyve* specific deletion in *Ptf1a-Cre* mice. F. Immunoblot analysis of pancreatic tissue from 12-week-old *Ptf1a-Cre; Pikfyve^+/+^*, *Ptf1a-Cre*; *Pikfyve^f/+^*, and *Ptf1a-Cre*; *Pikfyve^f/f^* mice showing changes in PIKfyve protein levels. Vinculin was used as a loading control. G. Densitometry analyses of immunoblot displayed in Fig. 1F. PIKfyve protein % was calculated by dividing the densitometry values for each PIKfyve band by the average value from the *Pikfyve^+/+^*group. (One-way ANOVA with Dunnett’s) H. Breeding design for the generation of KC *Pikfyve^+/+^*, KC *Pikfyve^f/+^*, KC *Pikfyve^f/f^* mice. I. Overall survival of KC *Pikfyve^+/+^*, KC *Pikfyve^f/+^*, KC *Pikfyve^f/f^* mice. Statistics were performed using a Gehan-Breslow-Wilcoxon test. J. Pancreas tissue weight normalized to total body weight from KC *Pikfyve^+/+^*, KC *Pikfyve^f/+^*, KC *Pikfyve^f/f^* or age-matched wild-type (WT) mice at 27 weeks of age. (One-way ANOVA with Dunnett’s) K. Percentage of pancreas occupied by normal tissue as determined by histological analyses in KC *Pikfyve^+/+^*, KC *Pikfyve^f/+^*, KC *Pikfyve^f/f^* mice at 27 weeks of age. (One-way ANOVA with Dunnett’s) L. Representative histological images showing H&E and CK19 staining on pancreatic tissue of KC *Pikfyve^+/+^*, KC *Pikfyve^f/+^*, KC *Pikfyve^f/f^* mice at 27 weeks of age. Scalebar = 5mm for the whole-pancreas images, 300μm for the zoomed-in images. M. Breeding design for the generation of KPC *Pikfyve^+/+^* and KPC *Pikfyve^f/f^* mice. N. *In situ Pikfyve* levels in KPC *Pikfyve^+/+^* and KPC *Pikfyve^f/f^*murine pancreas lesion vs normal tissue as determined by BaseScope. The KPC *Pikfyve^+/+^* scores used as a reference are the same as those used in Fig. 1B. The two cohorts were stained and analyzed in the same batch. (Multiple unpaired two-tailed t-test) O. Pancreas tissue weight normalized to total body weight from KPC *PIKfyve^+/+^* or KPC *PIKfyve^f/f^* mice at death. (Unpaired two-tailed t-test) P. Percentage of pancreas occupied by normal, pancreatic intraepithelial neoplasia (PanIN), or PDAC at death. (Two-way ANOVA). Q. Representative histology showing CK19 IHC and H&E staining of whole pancreatic tissue from KPC *PIKfyve^+/+^* and KPC *PIKfyve^f/f^* mice at 25 weeks. Scalebar = 5mm for the whole-pancreas images, 100μm for the high-magnification images.

We then sought to evaluate the effect of *Pikfyve* loss on PDAC development by crossing *Pikfyve^+/+^, Pikfyve^f/+^*, *or Pikfyve^f/f^* with the KC model (*Ptf1a-Cre; LSL-Kras^G12D/+^*) to assess pancreatic tumorigenesis **(Fig. 1H)**. We first confirmed a decrease in *Pikfyve* transcript in the pancreata of KC *Pikfyve^f/+^ and* KC *Pikfyve^f/f^* mice **(Extended Data Fig. 1D-E)**. In monitoring these cohorts of mice, we found that *Pikfyve* loss substantially extended the survival of mice harboring the KC genotype **(Fig. 1I)**. To determine whether this was correlated with a difference in pancreatic disease burden, we evaluated the pancreata of a separate cohort of mice and found that compared to pancreata of KC *Pikfyve*^+/+^ littermates, pancreata of KC *Pikfyve^f/+^* and KC *Pikfyve^f/f^* mice weighed less and were closer in weight to pancreata of wild-type mice at 27 weeks of age **(Fig. 1J, Extended Data Fig. 1F)**. Additionally, pancreata of KC mice with *Pikfyve* loss retained a higher degree of normal histological structures based on hematoxylin and eosin (H&E) staining or immunohistochemistry (IHC) staining for cytokeratin 19 (CK19) **(Fig. 1K-L)**. Consistent results were recapitulated at a later age of 40 weeks as well, both on macroscopic and microscopic evaluations **(Extended Data Fig. 1G-H)**.

We next evaluated the role of *Pikfyve* in the KPC model to assess the impact of PIKfyve on tumor progression **(Fig. 1M-N, Extended Data Fig. 1I)**. To do this, we harvested and analyzed 15 mice in the KPC *Pikfyve*^+/+^ cohort and 16 mice in the KPC *Pikfyve^f^*^/f^ cohort upon them reaching humane endpoints and found that the pancreata of the KPC *Pikfyve^f^*^/f^ mice weighed significantly less than those of KPC *Pikfyve*^+/+^ mice, relative to their total body weight **(Fig. 1O)**. To determine whether this effect was correlated with a decrease in disease onset or development, we performed histopathological analysis on these pancreata and observed that the pancreata of KPC *Pikfyve^f^*^/f^ mice displayed a significantly lesser degree of disease onset and progression compared to the pancreata of KPC *Pikfyve*^+/+^ mice **(Fig. 1P-Q)** at comparable ages **(Extended Data Fig. 1J).** Taken together, these data indicate that *Pikfyve* loss suppresses pancreatic cancer onset and progression in the KC and KPC models, respectively, while not affecting normal pancreatic tissue. Collectively, these studies with GEMMs suggest that PDAC has an elevated requirement for PIKfyve-driven processes.

### Pharmacological inhibition of PIKfyve suppresses PDAC development and growth

Given that genetic perturbation of *Pikfyve* attenuated PDAC development, we sought to evaluate whether pharmacological PIKfyve inhibition would elicit similar effects. We first confirmed that apilimod and ESK981, two PIKfyve inhibitors that have cleared phase 1 clinical trials^52,53^, bind to mouse PIKfyve protein using Cellular Thermal Shift Assay (CETSA) **(Fig. 2A)**. Given that apilimod is known to have poor *in vivo* pharmacokinetics^59^, we focused on ESK981 for subsequent *in vivo* experiments. To evaluate the impact of PIKfyve inhibition on PDAC development, we prophylactically treated a cohort of KPC mice aged to 6 weeks with ESK981 for 4 weeks **(Fig. 2B)**. At the endpoint of 10 weeks, we found that the weights of KPC pancreata treated with ESK981 were reduced to levels approaching that of wild-type pancreata **(Fig. 2C)**. On histopathological evaluation, ESK981-treated mice exhibited an increased retention of histopathologically unremarkable pancreatic tissue, including both acinar and endocrine components in normal physiological proportion, and relatively reduced PanIN and PDAC burden. These conclusions were based on exhaustive morphological evaluation by H&E, which were then broadly cross-validated by CK19 IHC staining **(Fig. 2D-E).**

**Figure 2:**
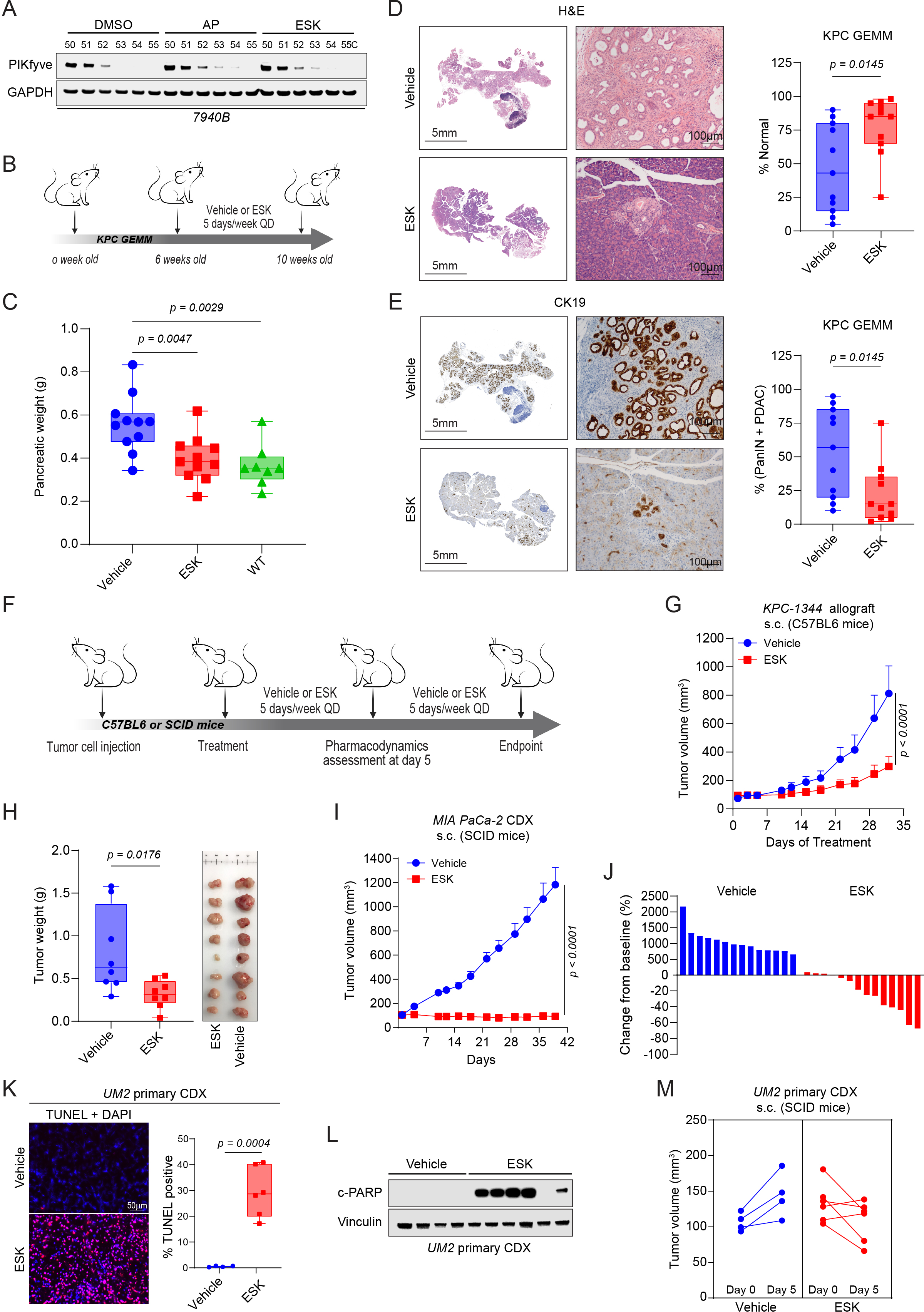
Pharmacological inhibition of PIKfyve blocks pancreatic cancer progression *in vivo*. A. Immunoblot analysis demonstrating stabilization of PIKfyve by apilimod (AP, 1000 nM) or ESK981 (ESK,1000 nM) in a cellular thermal shift assay (CETSA) employing the murine KPC cell line 7940B. B. Schematic of the *in vivo* study to assess prophylactic efficacy of vehicle or ESK981 (30 mg/kg) on KPC mice. C. Pancreatic tissue weight in vehicle- or ESK981 (30 mg/kg, QD, PO) -treated KPC mice in comparison with age-matched wild-type (WT) mice (right panel). (One-way ANOVA with Dunnett’s) D. Representative H&E staining of whole pancreatic tissue from vehicle and ESK981 treated mice (left). Quantification of histologically normal pancreatic tissue in vehicle or ESK981 treated mice (right). (Unpaired two-tailed t-test). GEMM: genetically engineered mouse model. Scalebar = 5mm for the whole-pancreas images, 100μm for the zoomed-in images. E. Representative CK19 IHC staining of whole pancreatic tissue from vehicle or ESK981 treated mice (left). Quantification of lesions (PanIN or PDAC) in vehicle or ESK981 treated mice (right). (Unpaired two-tailed t-test). Scalebar = 5mm for the whole-pancreas images, 100μm for the zoomed-in images. F. Schematic of *in vivo* efficacy studies utilizing cell-derived xenograft (CDX) or allograft models. ESK981 was dosed at 30 mg/kg per day (PO) in all studies. G. Tumor volumes of subcutaneous allograft model using KPC-derived KPC-1344 cells in response to vehicle or ESK981 in C57BL6 mice. Data plotted are mean tumor volumes + SEM (n=8 for each cohort). (Two-way ANOVA) H. Tumor weights (left) and images (right) of KPC-1344 model tumors at study endpoint. (Unpaired two-tailed t-test) I. Tumor volumes of subcutaneous CDX model using MIA PaCa-2 cells in response to vehicle or ESK981 in SCID mice. Data plotted are mean tumor volumes +SEM (n=14 for each cohort) (Two-way ANOVA). J. Waterfall plot displaying changes in tumor volume comparing endpoint to baseline in response to vehicle or ESK981 treatment. K. Left panel, representative images (one of three) of TUNEL staining from primary UM-2 CDX tumors after 5 days of treatment of vehicle or ESK981. Right panel, quantification of TUNEL positivity in indicated groups. Data plotted are from independent tumors and each represent the mean of 5 representative images per tumor. (Unpaired two-tailed t-test). Scalebar = 50μm L. Immunoblot analysis of primary UM2 CDX tumors after 5 days treatment of vehicle or ESK981 showing changes in apoptosis marker cleaved PARP (c-PARP). Vinculin was used as a loading control. M. Individual tumor volumes of a PDAC primary CDX UM-2 model before and after 5 days treatment of vehicle or ESK981.

Next, to determine the impact of PIKfyve inhibition on PDAC tumor growth, we employed *in vivo* allograft and xenograft models to test the efficacy of ESK981 **(Fig. 2F)**. In a KPC-derived subcutaneous syngeneic allograft, ESK981 therapy reduced tumor growth and weight at endpoint **(Fig. 2G-H)**. Similarly, ESK981 completely suppressed the growth of MIA PaCa-2 cell-derived xenograft (CDX) tumors **(Fig. 2I-J)**. To assess the impact of PIKfyve inhibition on non-KRAS-driven PDAC, we employed a BxPC-3 (*BRAF^V487-P492>A^*) CDX and showed that ESK981 still suppressed tumor growth and reduced tumor weight at endpoint **(Extended Data Fig. 2A-C)**. In both the MIA PaCa-2 and BxPC-3 CDX models, ESK981 treatment reduced the proliferation of these tumors based on Ki-67 staining **(Extended Data Fig. 2D)**. Further, ESK981 treatment induced substantial apoptosis in the MIA PaCa-2 as well as a UM2 (*KRAS^Q61L^*) primary CDX model, as shown by increased Terminal dUTP Nick End Labeling (TUNEL) staining and PARP cleavage **(Fig. 2K-L, Extended Data Fig. 2E)** after 5 days of treatment. We also observed regression in most of the tumors in the UM-2 primary CDX cohort upon ESK981 treatment **(Fig. 2M)**. Finally, we found that ESK981 treatment elicited similar effects on a *KRAS^G12V^*-driven T24 bladder CDX **(Extended Data Fig 2F-G)**. Taken together, these data show that PIKfyve inhibition decreases proliferation, induces apoptosis, and dramatically suppresses growth in both murine and human PDAC tumor models.

### PIKfyve perturbation suppresses autophagy and decreases PDAC cell proliferation

To determine the molecular effects of PIKfyve inhibition on PDAC cells, we employed a battery of methods to perturb PIKfyve. First, we employed CRISPR interference (CRISPRi), which decreased *PIKFYVE* transcript **(Extended Data Fig. 3A)** and protein levels **(Fig. 3A)** in the human PDAC cell lines MIA PaCa-2 and PANC-1 using two independent single guide RNAs (sgRNAs). *PIKFYVE* knockdown also increased the LC3A/B-II to LC3A/B-I ratio and increased p62 (SQSTM1) levels, suggesting an inhibition of autophagic flux **(Fig. 3A)**, consistent with data from previous reports^47,49^. Pharmacological inhibition of PIKfyve with apilimod (AP) or ESK981 (ESK) also showed similar effects in 7940B cells (murine KPC) and Panc 04.03 (human PDAC) cells as well as the UM-2 primary CDX tumors described in Figure 2 **(Fig. 3B, Extended Data Fig. 3B)**. As an orthogonal method to validate that PIKfyve inhibition decreases autophagic flux, we employed the GFP-LC3-RFP-LC3ΔG autophagic flux probe^60^. Treatment with apilimod, ESK981, or chloroquine (CQ) decreased basal autophagic flux, as well as autophagic flux induced by mTORC inhibition with torin-1 **(Fig. 3C, Extended Data Fig. 3C)**. Finally, as further confirmation of target specificity, we developed a second-generation proteolysis targeting chimera (PROTAC) degrader of PIKfyve, PIK5-33d, based on our previously described PIKfyve degrader^61^ **(Extended Data Fig. 3D)**. PIK5-33d (33d) potently degraded PIKfyve, and this phenocopied the autophagy inhibition phenotypes elicited by *PIKFYVE* knockdown or its enzymatic inhibition **(Extended Data Fig. 3E)**.

**Figure 3:**
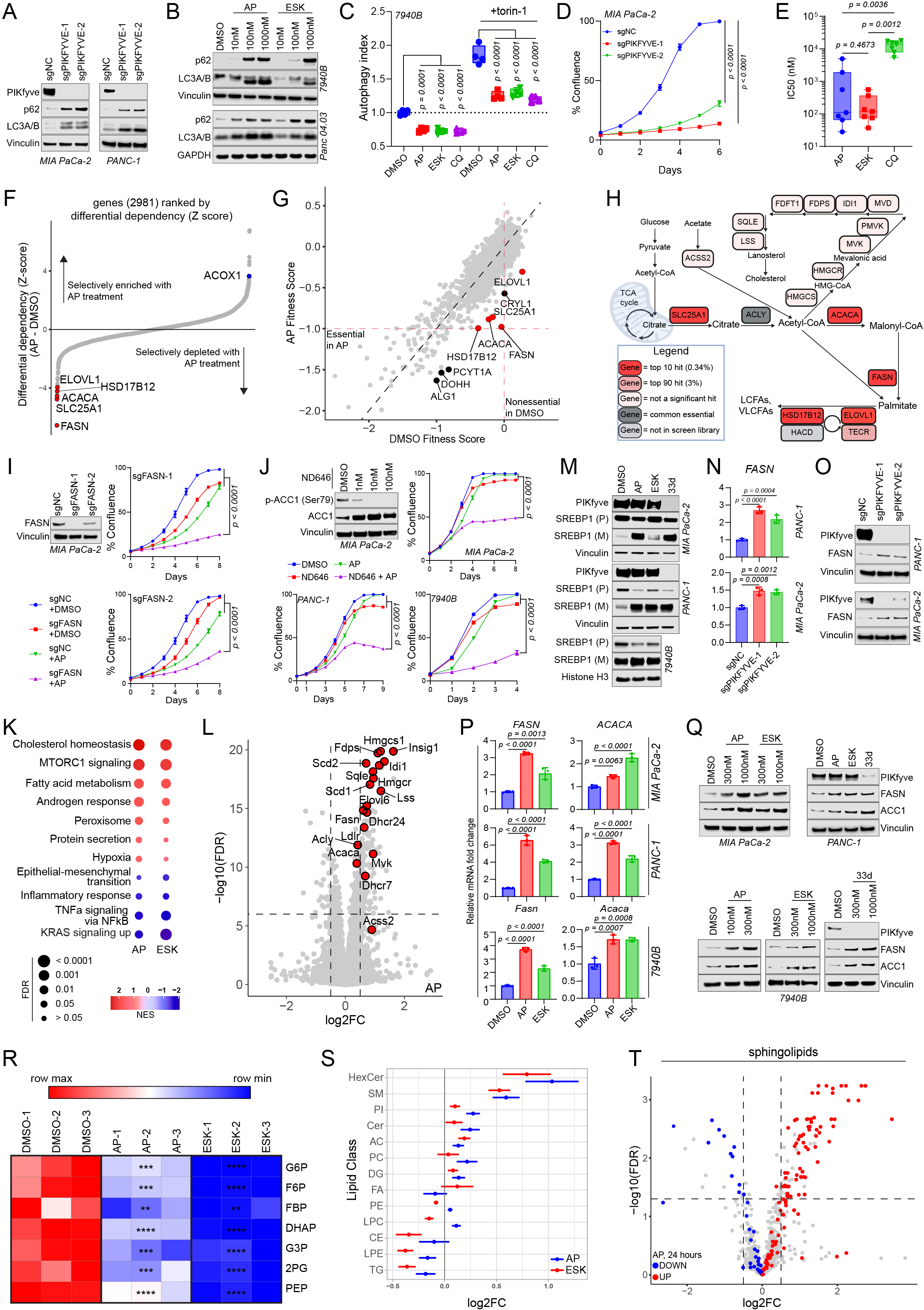
PIKfyve inhibition obligates PDAC cells to stimulate a lipogenic transcriptional and metabolic program. A. Immunoblot analysis of MIA PaCa-2 (left) and PANC-1 (right) cells upon CRISPRi-mediated knockdown of *PIKFYVE* with two independent sgRNAs (sgPIKFYVE-1 and sgPIKFYVE-2) or control (sgNC) showing changes in PIKfyve, p62 (SQSTM1), and LC3A/B. Vinculin was used as a loading control for all blots. This experiment was performed twice independently with similar results. B. Immunoblot analysis of known autophagy markers p62 (SQSTM1) and LC3A/B upon treatment with PIKfyve inhibitors apilimod (AP) or ESK981 (ESK) in KPC 7940B and Panc 04.03 cell lines. Vinculin or GAPDH were used as loading controls. The immunoblot using 7940B cells was performed twice independently with similar results. C. Tandem fluorescent autophagic flux reporter assay in 7940B cells after 24-hour treatment with apilimod (100 nM), ESK981 (1000 nM), and chloroquine (CQ, 50 μM) with or without mTORC1/mTORC2 inhibitor torin-1 (100 nM). Data shown are 4 biological replicates for each condition. (One-way ANOVA with Dunnett’s using indicated conditions as baseline). This represents one of three independent experiments. D. Confluence assay of MIA PaCa-2 cells upon CRISPRi-mediated knockdown of *PIKFYVE* (sgPIKFYVE) or control (sgNC). Data shown are mean +/- SEM (n=4 biological replicates) (Two-way ANOVA with Dunnet’s). This represents one of three independent experiments. E. Box-and-whisker plot displaying IC50s of apilimod, ESK981, and chloroquine in 7 human and mouse PDAC cell lines (specified in Extended Data Fig. 4 B-E). Statistics were performed using a Repeated Measures one-way ANOVA with Reisser-Greenhouse correction and with Tukey’s multiple comparisons test with individual variances computed for each comparison. F. Gene enrichment rank plot based differential sgRNA representation in apilimod-treated versus DMSO-treated endpoint populations of the CRISPR screen experiment. Lipid synthesis-related genes ranked at either extreme are highlighted. G. Scatter plot of gene fitness scores in apilimod-treated versus DMSO-treated endpoint conditions in metabolic CRISPR screen. Top 10 hits are labeled, and 5 lipid synthesis-related genes are highlighted. H. Metabolic map of fatty acid synthesis and elongation, and cholesterol homeostasis. Red indicates top 10 hit in the CRISPR screen; pink indicates top 90 (3%) hit; light pink indicates the gene was not a top 3% hit; dark grey indicates the gene is universally essential; light grey indicates the gene was not included in the CRISPR screen library. I. Immunoblot of MIA PaCa-2 upon CRISPRi-mediated knockdown of *FASN* in cells and corresponding confluence assays assessing the sensitivity of *FASN* knockdown (sgFASN) or control (sgNC) cells to apilimod (100 nM). Vinculin was used as a loading control for the immunoblot. Confluence assay data shown are mean +/- SEM (n=4) from one of two independent experiments. Statistics were performed using an F statistics test based on a two-way ANOVA with Dunnett’s multiple comparisons test with the sgFASN + apilimod condition set as a baseline. J. Immunoblot analysis assessing the phosphorylation status of ACC1 (p-ACC1) in MIA PaCa-2 cells upon ND646 (ACC inhibitor) treatment and corresponding confluence assays assessing the sensitivity of MIA PaCa-2, PANC-1, and 7940B cells to apilimod, ND646, or both. Vinculin was used as a loading control for the immunoblot. Confluence assay data shown are mean +/- SEM (n=4). Statistics were performed using a two-way ANOVA with Dunnett’s multiple comparisons test with the ND646+apilimod condition set as a baseline. Concentrations used for apilimod are: 100 nM for MIA PaCa-2 and 50 nM for PANC-1 and 7940B. Concentrations used for ND646 are: 100 nM for MIA PaCa-2 and 1000 nM for PANC-1 and 7940B. The confluence assays were performed three times independently with similar results. The DMSO and ND646 conditions for MIA PaCa-2 and PANC-1 are also utilized as controls in Extended Data Fig. 6F and G as these data were generated from the same experiment. K. Pathway enrichment analysis of RNA-seq performed on 7940B treated with either apilimod (25 nM) or ESK981 (250 nM) for 8 hours. Dot sizes are inversely proportional to false discovery rate (FDR). The color scheme is reflective of the normalized enrichment score (NES). L. Volcano plot using RNA-seq analysis on 7940B cells treated with apilimod (25 nM) for 8 hours highlighting SREBP-1 target genes. Vertical dashed lines indicate log2 fold change = +/- 0.5). Horizontal dashed line indicates FDR=10^-6^. M. Immunoblot showing PIKfyve, premature SREBP1 (SREBP1 (P)), and mature SREBP1 (SREBP1 (M)) in MIA PaCa-2, PANC-1, and 7940B cells upon treatment with PIKfyve inhibitors or degrader PIK5-33d (33d) for 8 hours. Vinculin or histone H3 were used as loading controls. The drug doses used were as follows: MIA-PaCa-2 and PANC-1: apilimod=300 nM, ESK981=1000 nM, PIK5-33d=1000 nM; 7940B: apilimod=100 nM, ESK981=1000 nM, PIK5-33d=1000 nM. This data is representative of two independent experiments. N. Quantitative-PCR (qPCR) of MIA PaCa-2 and PANC-1 showing changes in RNA levels of *FASN* upon CRISPRi-mediated knockdown of *PIKFYVE* using two independent sgRNAs targeting *PIKFYVE* compared to control. Data plotted are technical triplicates from one of three independent experiments. (One-way ANOVA with Dunnett’s). O. Immunoblot analysis of MIA PaCa-2 and PANC-1 showing changes in protein levels of FASN upon CRISPRi-mediated knockdown of *PIKFYVE* using two independent sgRNAs targeting *PIKFYVE* relative to control. Vinculin was used as a loading control. These data are representative of two independent experiments. P. qPCR of MIA PaCa-2, PANC-1, and 7940B showing changes in RNA levels of labeled genes upon treatment with PIKfyve inhibitors for 8 hours. The drug doses used were as follows: MIA PaCa-2 and PANC-1: apilimod = 300 nM, ESK981 = 1000 nM. 7940B: apilimod = 100 nM, ESK981 = 1000 nM. Data plotted are technical triplicates (One-way ANOVA with Dunnett’s). These experiments were performed three independent times each with similar results. Q. Immunoblot analysis of MIA PaCa-2, PANC-1, and 7940B showing changes in protein levels of labeled genes upon treatment with PIKfyve inhibitors for 24 hours. Vinculin was used as a loading control. The drug doses used were indicated on figure or as follows for PANC-1: apilimod = 300 nM, ESK981 = 1000 nM, PIK5-33d = 1000 nM. These data are representative of two independent experiments each. R. Heatmap of glycolytic metabolite abundance in 7940B cells treated with DMSO, apilimod (100 nM), or ESK981 (1000 nM) for 8 hours. G6P = glucose 6-phosphate; F6P = fructose 6-phosphate; FBP = fructose 1,6-bisphosphate; DHAP = dihydroxyacetone phosphate; G3P = glyceraldehyde 3-phosphate; 2PG = 2 phosphoglycerate; PEP = phosphoenolpyruvate. * indicates p < 0.05, ** indicates p < 0.01, *** indicates p < 0.001, **** indicates p < 0.0001. (One-way ANOVA with Dunnett’s) S. Forest plot indicating changes in lipid class abundance in 7940B cells upon treatment with DMSO, apilimod (100 nM) or ESK981 (1000 nM) for 24 hours. HexCer = hexosylceramide; SM = sphingomyelin; PI= phosphatidylinositol; Cer = ceramide; AC= acylcarnitine; PC= phosphatidylcholine; DG = diacylglyceride; FA = fatty acid; PE = phosphatidylethanolamine; LPC = lysophosphatidylcholine; CE = cholesteryl ester; LPE = lipophosphatidylethanolamine; TG = triacylglyceride. Effect sizes are in log2 scale of lipid abundance estimated from separate linear model for each treatment (apilimod or ESK981) compared to DMSO, adjusting for lipid classes with random intercept. T. Volcano plot of lipidomics analysis on 7940B cells treated with apilimod for 24 hours, plotting log2 fold change compared to DMSO highlighted changes in sphingolipid classes (HexCer, SM, Cer). Red highlights indicate upregulated sphingolipids; blue highlights indicate downregulated sphingolipids. Vertical dashed lines indicate log2 fold change = +/- 0.5). Horizontal dashed line indicates *p = 0.05.* (unpaired two-tailed t-test)

Consistent with previous work, *PIKFYVE* knockdown cells revealed a lysosomal vacuolization phenotype, which was also evident within four hours of PIKfyve inhibitor or degrader treatment **(Extended Data Fig. 3F, G)**^47,49^. Importantly, consistent with our tumor studies, PIKfyve perturbation through *PIKFYVE* knockdown substantially slowed the growth of PDAC cells **(Fig. 3D, Extended Data Fig. 4A)**, and PIKfyve inhibition decreased PDAC cell viability with half-maximal inhibitory concentrations (IC_50_) in the nanomolar ranges for most cell lines **(Extended Data Fig. 4B-C)**. Lysosome inhibition by chloroquine treatment also decreased PDAC cell viability **(Extended Data Fig. 4D)**; however, the IC_50_ values were much higher for chloroquine than apilimod or ESK981 in the same PDAC cell lines (**Fig. 3E, Extended Data Fig. 4E)**. Taken together, these data illustrate that PIKfyve plays a crucial role in regulating autophagy, lysosomal homeostasis, and, ultimately, cell proliferation in PDAC.

PDAC is known to utilize autophagy and lysosomal processes to promote iron homeostasis and allow for mitochondrial respiration^23,24,55^; therefore, we investigated whether PIKfyve inhibition decreased PDAC cell proliferation through a similar mechanism. PIKfyve inhibition stabilized HIF1α upon eight hours of treatment **(Extended Data Fig. 5A)**, consistent with the effect of iron deprivation due to disrupting autophagy. However, PIKfyve inhibition did not decrease basal oxygen consumption rate (OCR) in 7940B or Panc 04.03 cells, contrasting the activity of chloroquine (CQ) and bafilomycin A1 (BAF), the other autophagy and lysosomal inhibitors tested **(Extended Fig. 5B)**. Consistent with this, PIKfyve inhibition had no impact on OCR through 24 hours of treatment, compared to CQ and BAF, which significantly decreased OCR in 7940B cells starting from eight hours **(Extended Data Fig. 5C)**. To further confirm that PIKfyve inhibition does not decrease PDAC cell proliferation through disrupting iron homeostasis, we attempted to rescue PDAC cells from PIKfyve inhibition using ferric ammonium citrate (FAC). While the antiproliferative effects of BAF were drastically attenuated by addition of FAC, we did not see a similar effect with PIKfyve inhibitors **(Extended Data Fig. 5D-F)**. Overall, these data suggest that autophagy and lysosomal perturbation through PIKfyve inhibition does not decrease PDAC proliferation by disrupting iron homeostasis and mitochondrial respiration but, rather, occurs through a distinct mechanism.

### PIKfyve inhibition creates a synthetic lethality of *de novo* lipid synthesis in PDAC cells

To assess the functionally relevant metabolic roles of PIKfyve in PDAC in an unbiased manner, we employed a metabolism-focused CRISPR screen in MIA PaCa-2 cells treated with apilimod **(Extended Data Fig. 6A)**. This screen accurately discriminated against core essential and non-essential genes, validating its biological relevance and consistency **(Extended Data Fig. 6B)**. Interestingly, the most significantly depleted sgRNAs targeted genes core to the *de novo* fatty acid synthesis and elongation pathways, namely *FASN*, *ACACA*, *SLC25A1*, *ELOVL1*, and *HSD17B12* **(Fig. 3F-G, Supplementary Table 1)**. In contrast, *ACOX1*, which completes the first step of lipid beta-oxidation, was the target of some of the most significantly enriched sgRNAs in the screen **(Fig. 3F)**. Additionally, no cholesterol-specific genes were among the significant hits, suggesting that *de novo* fatty acid synthesis was a specific, functionally relevant synthetic essentiality of MIA PaCa-2 cells upon PIKfyve inhibition **(Supplementary Table 1, Fig. 3H)**. To validate this screen, we employed CRISPRi-mediated knockdown of *FASN* in MIA PaCa-2 cells and found that *FASN* knockdown with two independent sgRNAs **(Extended Data Fig. 6C)** indeed sensitized cells to apilimod **(Fig. 3I)** and the PIKfyve degrader PIK5-33d (33d) **(Extended Data Fig. 6D)**. As an orthogonal validation, we utilized ND646, which is an inhibitor of ACC1 (protein name of *ACACA*). After confirming on-target effects of ND646 using immunoblots **(Fig. 3J, Extended Data Fig. 6E)**, we found that ND646 similarly sensitized PDAC cells to apilimod **(Fig. 3J)**, ESK981 **(Extended Data Fig. 6F)**, and PIK5-33d **(Extended Data Fig. 6G)** using MIA PaCa-2, PANC-1, and 7940B cell lines. These data suggest that upon PIKfyve inhibition, PDAC cells become reliant on the *de novo* fatty acid synthesis pathway to proliferate.

### PIKfyve inhibition promotes the upregulation of *de novo* lipid synthesis in PDAC cells

Given that PIKfyve inhibition obligates PDAC cells to maintain expression and function of the *de novo* fatty acid synthesis pathway, we next assessed whether PIKfyve perturbation caused upregulation of this pathway. Utilizing RNA-seq in 7940B cells, we determined that eight-hour treatment of apilimod or ESK981 induced remarkably concordant gene expression changes **(Extended Data Fig. 7A)**, and the most upregulated pathways were related to cholesterol homeostasis, MTORC1 signaling, and fatty acid metabolism **(Fig. 3K, Extended Data 7B)**. Additionally, most of the top upregulated genes were targets of transcription factor sterol regulatory element binding transcription factor 1 (SREBP1), a key regulator of lipogenesis^62^ **(Fig 3L, Extended Data Fig. 7C)**. Accordingly, we confirmed that eight hours of PIKfyve inhibition or degradation activated SREBP1 by post-translational cleavage **(Fig. 3M)**. Importantly, *FASN* was upregulated upon *PIKFYVE* knockdown **(Fig. 3N-O)**, and both FASN and ACC1 were upregulated upon PIKfyve inhibition at the transcript **(Fig. 3P)** and protein levels **(Fig. 3Q)**. Taken together, these results illustrate that PDAC cells upregulate a lipogenic transcriptional program in response to PIKfyve inhibition.

To determine whether the lipogenic transcriptional program translated to a metabolic phenotype, we employed metabolic analyses on 7940B cells. PIKfyve inhibition, using apilimod or ESK981 treatment, induced a similar metabolic landscape (**Extended Data Fig. 8A)** featuring a decrease in citrate at three hours of treatment **(Extended Data Fig. 8B)**. At eight hours, the citrate level recovered to comparable levels to the DMSO condition **(Extended Data Fig. 8B)**; however, this was associated with a dramatic decrease of upstream glycolytic metabolites **(Fig. 3R, Extended Data Fig. 8C)**. Given that the citrate transporter SLC25A1 was also a top hit in the CRISPR screen **(Fig. 3H)**, we hypothesized that the glycolytic metabolites were being utilized to generate citrate that was then shunted into *de novo* lipid synthesis. To verify this, we performed targeted lipidomics and found that PIKfyve inhibition, whether through apilimod or ESK981 treatment, induced significant changes in the cellular lipid landscape in 7940B cells **(Extended Data Fig. 8D)**. We then grouped the lipid species into their respective classes and determined that hexosylceramides (HexCer), sphingomyelin (SM), and ceramide (Cer) were three of the top four upregulated lipid classes **(Fig. 3S)**. These classes, all sphingolipids, contained the majority of the significantly upregulated lipid species **(Fig. 3T, Extended Data Fig. 8E)**. These data suggest that upon PIKfyve loss of function, PDAC cells are forced to increase *de novo* lipid synthesis and accumulate sphingolipids as a survival mechanism.

### KRAS-MAPK regulates *de novo* lipid biosynthesis in PDAC

To identify avenues to possibly leverage the synthetic lethality of PIKfyve and *de novo* fatty acid synthesis, we sought to determine drivers of *FASN* and *ACACA* transcription in PDAC. KRAS is known to be a core driver of metabolic homeostasis in PDAC through MAPK signaling^63^; thus, we determined whether KRAS-MAPK signaling drove *FASN* and *ACACA* expression. Employing an inducible *Kras*^G12D^ cell line, iKras 9805 (iKras)^64^, we found that doxycycline withdrawal (Kras OFF) decreased *Fasn* and *Acaca* expression at the transcript **(Extended Data. Fig. 9A)** and protein level **(Fig. 4A)**. Further, eight-hour treatment with MRTX1133 (MRTX, KRAS^G12D^ inhibitor), AMG510 (AMG, KRAS^G12C^ inhibitor), or trametinib (MEK inhibitor) decreased transcription of *FASN* and *ACACA* **(Extended Data Fig. 9B)** in PDAC cell lines with the relevant KRAS mutation. This was reflected by a decrease in protein level after 48 hours of treatment **(Fig. 4B)**. These data are concordant with previously published RNA-seq data suggesting that MRTX1133 treatment decreases *FASN* and *ACACA* transcripts in AsPC1 cells *in vitro* **(Extended Fig. 9C)** and *in vivo* **(Extended Fig. 9D)**^43^. All together, these data illustrate that KRAS-MAPK signaling regulates *FASN* and *ACACA* expression in PDAC.

**Figure 4:**
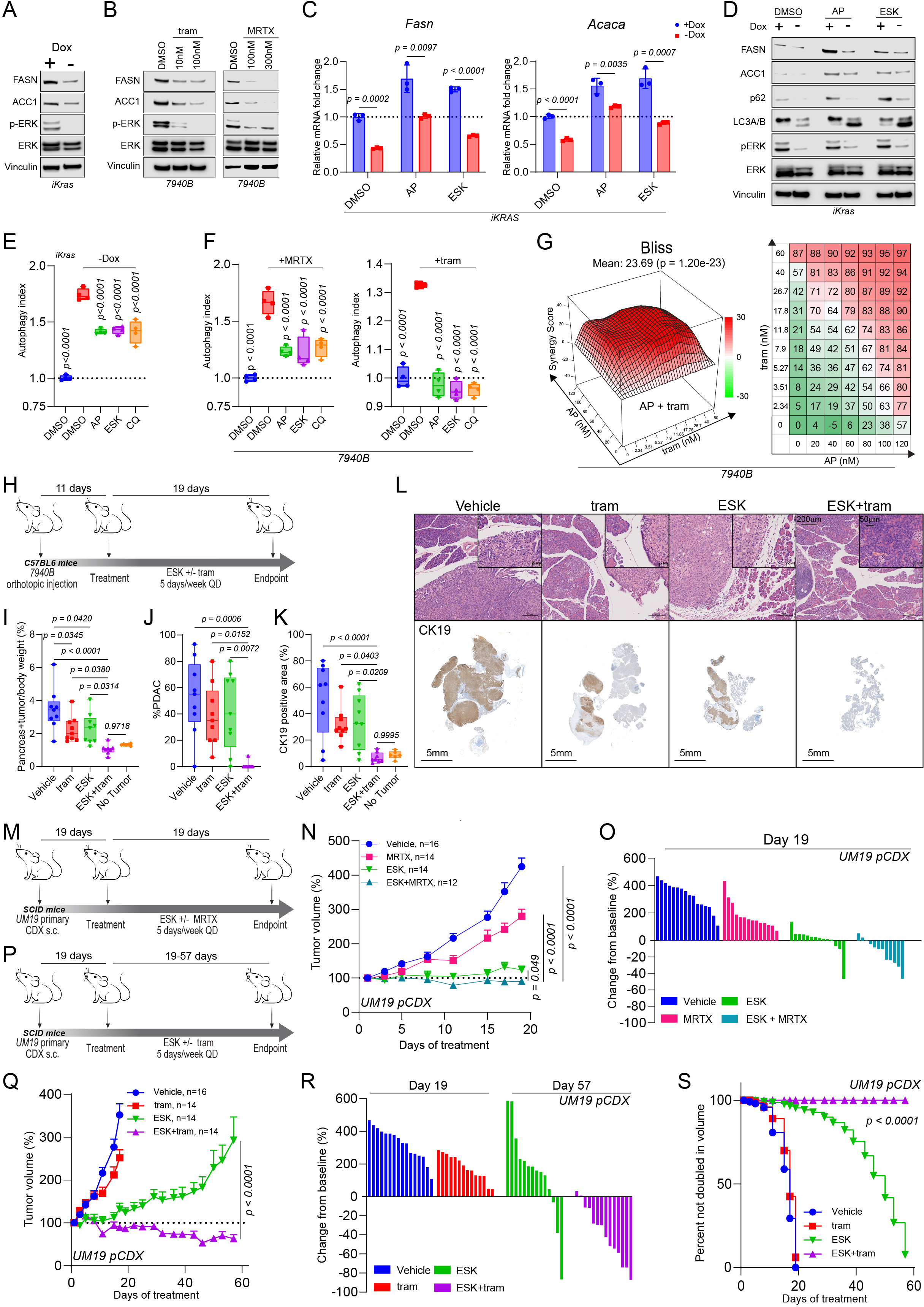
Dual KRAS-MAPK and PIKfyve inhibition results in metabolic crises and synergistic growth suppression in PDAC. A. Immunoblot analysis of iKRAS 9805 cells showing changes in protein levels of FASN and ACC1 upon presence or absence of doxycycline for 72 hours. Phospho-ERK and ERK were used to validate KRAS-MAPK signal inhibition. Vinculin was used as a loading control. This is representative of two independent experiments. B. Immunoblot analysis of 7940B cells treated with MEK inhibitor trametinib (tram) or KRAS^G12D^ inhibitor MRTX1133 (MRTX) for 48 hours at the indicated concentrations showing changes in protein levels of FASN and ACACA. Phospho-ERK and ERK were used to validate on-target effects on KRAS-MAPK signaling. Vinculin was used as a loading control. MRTX1133 and DMSO were refreshed every 12 hours in experiments involving MRTX1133. These experiments were each performed independently twice with similar results. C. qPCR of iKRAS 9805 cells showing changes in mRNA levels of *Fasn* (left) and *Acaca* (right) upon 48-hour incubation with or without doxycycline (Dox) and subsequent 8-hour treatment with apilimod (50 nM), ESK981 (300 nM), or DMSO. Data shown are technical replicates (multiple unpaired two-tailed t-tests). This experiment was performed independently three times each. D. Immunoblot analysis of iKRAS 9805 cells showing changes in protein levels of FASN, ACC1, p62, and LC3A/B upon 48-hour incubation with or without doxycycline (Dox) and subsequent 24-hour treatment with apilimod (50 nM), ESK981 (300 nM), or DMSO. p-ERK and ERK were assessed to validate KRAS OFF. Vinculin was used as a loading control. This is representative of two independent experiments. E. Tandem fluorescent reporter assay on iKRAS 9805 cells showing changed autophagic flux upon doxycycline withdrawal for 24 hours and subsequent treatment with apilimod (100 nM), ESK981 (1000 nM), or chloroquine (10 μM) for 24 hours. (One way ANOVA with Dunnett’s). This data is representative of three independent experiments each. F. Tandem fluorescent reporter assay on 7940B cells showing changed autophagic flux upon 4-hour pretreatment with apilimod (100 nM), ESK981 (1000 nM), or chloroquine (50 μM) and subsequent treatment with MRTX1133 (300 nM) or trametinib (25 nM) for 24 hours (One-way ANOVA with Dunnett’s). This data is representative of three independent experiments each. G. 3D synergy and corresponding heatmap plots in 7940B cells treated with apilimod and trametinib. Red peaks in the 3D plots indicate synergism, and the overall average synergy score is listed above. The heatmap plots the decrease in viability in 7940B upon treatment with each single agent or combination across indicated doses of each inhibitor. This experiment was performed three independent times with similar results. H. Schematic outlining syngeneic orthotopic model of 7940B for C57BL/6 mice assessing *in vivo* efficacy of ESK981 (ESK, 30 mg/kg, QD, PO), trametinib (tram, 1 mg/kg QD, PO), or ESK981 and trametinib (ESK + tram). I. Endpoint pancreas + tumor weight normalized to total body weight. Pancreata of 6 age-matched non-tumor bearing C57BL/6 mice were used as references. (One-way ANOVA with Tukey’s) J. Quantification of proportion of PDAC in H&E section from each tumor of the 7940B syngeneic orthotopic model. (One-way ANOVA with Tukey’s) K. Quantification of CK19 positive area compared to hematoxylin counterstain on a section from each tumor of the 7940B syngeneic orthotopic model. (One-way ANOVA with Tukey’s) L. Representative images of H&E and CK19 IHC staining of one tumor from each treatment arm from 7940B syngeneic orthotopic model. Scalebar = 200μm for the zoomed-out H&E images; 50μm for the zoomed-in H&E images; and 5mm for the whole-pancreas CK19 IHC images. M. Schematic outlining efficacy study using subcutaneous model of UM-19 primary cell-derived xenograft (CDX) treated with vehicle, MRTX1133 (MRTX, 30 mg/kg, QD, IP), ESK981 (ESK 30 mg/kg, QD, PO), or ESK981 + MRTX1133 (ESK + MRTX). N. Tumor volumes as a percentage (+SEM) of the initial volume measured by calipers over treatment course of the UM19 primary CDX (pCDX) model treated with MRTX1133 +/- ESK981. Statistics were performed using a two-way ANOVA with Dunnett’s, with the combination group used as the reference. O. Waterfall plot displaying change in tumor volume at treatment end point (day 19) compared to baseline of the UM19 pCDX model treated with MRTX1133 +/- ESK981. P. Schematic outlining efficacy study using subcutaneous model of UM19 pCDX treated with vehicle, trametinib (tram, 1 mg/kg, QD, PO), trametinib (tram 1 mg/kg QD, PO), ESK981 (ESK 30 mg/kg, QD, PO), or ESK981 + trametinib (ESK + tram). Q. Tumor volumes as a percentage (+SEM) of the initial volume measured by calipers over treatment course of the UM19 pCDX model treated with trametinib +/- ESK981. Statistics were performed using a two-way ANOVA with Dunnett’s, with the combination group used as the reference. The tumors in the vehicle- and ESK981-treated groups were the same tumors shown in Fig. 4O. R. Waterfall plot displaying change in tumor volume at treatment end point compared to baseline of the UM19 CDX model treated with trametinib +/- ESK981. The endpoint displayed of the vehicle and trametinib arms are day 19. The endpoint displayed of the ESK981 and ESK981 +/- trametinib group are day 57. The tumors in the vehicle- and ESK981- treated groups were the same tumors shown in Fig. 4O. S. Kaplan–Meier estimates of time to tumor doubling (Gehan-Breslow-Wilcoxon test).

To directly assess the effects of dual inhibition of PIKfyve and KRAS on FASN and ACC1, we treated iKras cells with PIKfyve inhibitors after incubation with or without doxycycline. While PIKfyve inhibition increased the transcription of *Fasn and Acaca*, concurrent *Kras* OFF and PIKfyve inhibition led to lesser increase of *Fasn* and *Acaca* transcript levels compared to baseline **(Fig. 4C)**. In a similar fashion, PIKfyve inhibition increased the protein levels of FASN and ACC1, while concurrent *Kras* OFF and PIKfyve inhibition led to a lesser increase in FASN and ACC1 protein levels compared to baseline **(Fig. 4D)**. This data suggests that KRAS-MAPK inhibition blocks expression of *FASN* and *ACACA*, synthetically critical genes in PDAC upon PIKfyve inhibition.

### Concurrent perturbation of PIKfyve and KRAS-MAPK creates metabolic conflict of autophagy regulation

Important recent studies revealed that PDAC cells upregulate and depend on autophagy to maintain metabolic homeostasis upon KRAS-MAPK signaling inhibition^31-33^. Utilizing the autophagic flux probe, we confirmed that PDAC cells upregulate autophagy upon acute mutant KRAS^G12D^ inhibition with MRTX-1133 **(Extended Data Fig. 9E)**. Knowing that PIKfyve inhibition disrupts autophagic flux, we sought to determine whether PIKfyve inhibition could also leverage this induced metabolic dependency upon KRAS perturbation. Indeed, *Kras* OFF induced an increase in the LC3-II to LC3-I ratio, and this ratio was maintained when PIKfyve inhibitors were added **(Fig. 4D)**. Additionally, p62 was decreased upon *Kras* OFF, increased upon PIKfyve inhibition, and less dramatically changed with both *Kras* OFF and PIKfyve inhibition **(Fig. 4D)**. This suggests that PIKfyve inhibition and *Kras* OFF exert opposing effects on autophagic flux. We validated this using the autophagic flux probe assay in iKras cells, which showed an increase in autophagic flux upon *Kras* OFF that was attenuated with PIKfyve inhibition or chloroquine treatment **(Fig. 4E)**. Pharmacological inhibition of KRAS-MAPK using MRTX1133 or trametinib also induced autophagic flux that was blocked upon PIKfyve inhibition in 7940B **(Fig. 4F)** and Panc 04.03 cells **(Extended Data Fig. 9F)**. Altogether, this suggests that concurrent PIKfyve and KRAS-MAPK inhibition drives PDAC into a state of metabolic conflict regarding its regulation of autophagic flux.

### Dual inhibition of PIKfyve and KRAS-MAPK synergistically suppresses PDAC growth

We next sought to assess whether the metabolic crises elicited by simultaneous inhibition of PIKfyve and KRAS-MAPK could be utilized to inhibit PDAC cell proliferation. Synergy assays confirmed that any combination of PIKfyve inhibition, using apilimod or ESK981, and KRAS-MAPK inhibition, using MRTX1133 or trametinib, resulted in striking synergistic effects, decreasing PDAC cell proliferation and viability **(Fig. 4G, Extended Data Fig. 10A-C).**

To determine the efficacy of combining PIKfyve and KRAS-MAPK inhibitors as a therapeutic strategy for PDAC, we utilized a syngeneic orthotopic preclinical model **(Fig. 4H)**. Importantly, treatment with ESK981 and/or trametinib did not impact mouse body weight throughout the treatment course **(Extended Data Fig. 11A)**. Upon endpoint analysis, we did not observe any gross evidence of tumor burden in any mice treated with the combination of ESK981 and trametinib. To ensure that we accounted for microscopic tumor burden, we weighed the tumors and pancreata together for each of the mice. Upon completing this analysis, we observed that the mice treated with the combination had significantly lighter pancreata **(Extended Data Fig. 11B)**, comparable to those found in age-matched, non-tumor bearing mice, while the individual treatments had more modest effects compared to vehicle-treated mice **(Fig. 4I)**. Histopathological evaluation with H&E and CK19 corroborated this, revealing no evidence of PDAC in seven out of eight mice treated with both ESK981 and trametinib, while either treatment alone exhibited only marginal effects **(Fig. 4J-L)**. Taken together, these data illustrate that combination therapy of a PIKfyve inhibitor and MEK inhibitor eliminated tumor burden in an immunocompetent orthotopic PDAC model **(Extended Data Fig. 11C)**.

Next, we further tested this therapeutic strategy in a human PDAC model utilizing UM19, a primary *KRAS*^G12D^ PDAC CDX (pCDX) **(Fig. 4M)**. Combination treatment of ESK981 and MRTX1133 significantly improved the efficacy of either treatment alone throughout the treatment duration **(Fig. 4N)**. At endpoint, the combination induced regression in nearly all tumors, while each individual treatment had more modest effects **(Fig. 4O)**. Further, combining ESK981 and trametinib **(Fig. 4P)** induced substantial and durable regression in nearly all tumors, even when the tumors were able to adapt and outgrow ESK981 or trametinib therapy alone **(Fig. 4Q)**. At endpoint, most of the tumors treated with the combination were still smaller than their original size, some essentially undetectable **(Fig. 4R)**. Ultimately, the combination prevented any tumor from doubling throughout the duration of the experiment, while nearly all tumors from the other treatment groups doubled or more in size **(Fig. 4S)**.

In sum, these data demonstrate that KRAS-MAPK inhibition creates a synthetic vulnerability to PIKfyve inhibition *in vitro* and *in vivo*. Unlike previous efforts to target autophagy in PDAC, ESK981 has vastly superior pharmacological properties^52^. Further, the arrival of KRAS inhibitors provides exciting context to explore this combination in the clinic, noting the safety profile of the combination in our studies.

## Discussion

Targeting lysosome function and the autophagic pathway as a therapeutic strategy has shown promise preclinically, given the known metabolic vulnerabilities of PDAC^18,46^. Further enhancing this concept was the important finding that PDAC utilizes autophagy to compensate for KRAS-MAPK inhibition^31-33^. However, hydroxychloroquine (HCQ), the only clinical-grade compound available to target these pathways, has had limited efficacy^28,29^. In addition, HCQ (and its predecessor chloroquine, CQ) does not have a definitive molecular target, making it suboptimal for systematic pharmacological development^65^. In this study, we nominated PIKfyve, a lipid kinase known for its important roles in lysosomal function^7^, as a druggable target to leverage PDAC’s metabolic vulnerabilities of nutrient scavenging and recycling through the lysosome. In our studies, we discovered that *PIKFYVE* is expressed at a higher level in PDAC compared to normal pancreas in both human patient and murine PDAC samples, suggesting that PDAC cells have an increased need for PIKfyve activity compared to healthy pancreatic cells. Further, we showed that *Pikfyve* knockout or inhibition with the phase 1-cleared inhibitor ESK981 substantially decreased tumor development and growth in murine and human *in vivo* models, suggesting that PIKfyve is essential for PDAC development and growth. Taken together, these data highlight PIKfyve as the first gene involved in autophagy/lysosome function for which there exists both genetic and clinically relevant pharmacologic evidence of its viability as a therapeutic target in PDAC. This promising preclinical data has been used to position a multi-center phase 2 clinical trial (NCT05988918) assessing the efficacy of ESK981 on solid tumors, including PDAC.

Though lysosomal processes and autophagy have long been identified as metabolic targets for PDAC, the exact roles they play in PDAC metabolic homeostasis remain unclear. Recent work using a metabolism-focused CRISPR screen in an acute T cell leukemia line with the V-ATPase inhibitor bafilomycin and ammonia demonstrated that lysosomes serve a crucial role in maintaining iron homeostasis^55^. Multiple reports have independently verified and expanded on this concept in various models, including PDAC^25,66^. In our studies, we approached autophagy and lysosomal inhibition using a different well-defined target (PIKfyve) and an exquisitely specific inhibitor (apilimod^50^). Applying a similar CRISPR screening library in MIA PaCa-2 cells with apilimod, we were surprised to find that five out of the top ten genes that were scored as selectively essential were core to the fatty acid synthesis and elongation pathway, such as *FASN* and *ACACA* **(Fig. 3F-H)**. The RNA-seq experiment further highlighted lipid metabolism as the most dramatically affected gene signatures upon PIKfyve inhibition **(Fig. 3K)**. Taken in context, our data raises the possibility that specific methods of inhibiting lysosomal processes may have differential effects on various aspects of PDAC cell metabolism.

While we believe that this study is the first to identify a relationship of synthetic lethality between PIKfyve and fatty acid synthesis in PDAC, PIKfyve was recently shown to play a role in lipid metabolism through its role in lysosome function^67^. In this study, the authors inhibited *de novo* fatty acid synthesis and found that cells undergo increased phospholipid turnover in a lysosome- and PIKfyve-dependent process. In a converse manner, our study identified that PIKfyve inhibition stimulated *de novo* fatty acid synthesis and elongation. Together, our studies provide independent and complementary evidence that PIKfyve plays a crucial role in maintaining lipid homeostasis in coordination with *de novo* fatty acid synthesis, suggesting that disruption of one arm increases cells’ dependence on the other. Thus, a logical implication made by our studies would be that simultaneous perturbation of both arms would lead to catastrophic metabolic dysregulation.

The therapeutic strategy of inhibiting both KRAS-MAPK and autophagy has gained considerable recent attention, including being the subject of recent clinical trials (NCT04386057, NCT04132505)^31-33^. In the studies describing this relationship, the authors identified that PDAC utilizes autophagy as an adaptive and protective mechanism to maintain metabolic homeostasis upon KRAS-MAPK inhibition. Knowing PIKfyve’s role in autophagy, this alone positions PIKfyve inhibitors as alternatives to CQ to pair with KRAS-MAPK inhibitors for PDAC therapy **(Fig. 4E-F)**. However, through this study, we believe we have discovered a an additional mechanistically distinct rationale for dual inhibition of PIKfyve and KRAS-MAPK. Through the metabolic CRISPR screen, we identified that PIKfyve and fatty acid synthesis have a robust relationship of synthetic lethality **(Fig. 3F-H)**. In a search for translatable methods for leveraging this novel relationship, we hypothesized that KRAS-MAPK would drive fatty acid synthesis. Indeed, we found that KRAS-MAPK perturbation transcriptionally downregulated key fatty acid synthesis genes *FASN* and *ACACA* **(Fig. 4A-D)**. This positions KRAS-MAPK inhibitors as promising combinatorial partners with PIKfyve inhibitors for PDAC therapy. We assessed this combination in both *in vitro* and *in vivo* preclinical models and found in each case that the combination exhibited dramatically more potent effects than the individual therapies, in some cases even eliminating tumor burden.

In summary, we nominate PIKfyve as a preeminent therapeutic target to disrupt PDAC lysosomal function, a unique metabolic dependency of PDAC. Supporting this, we showed that PIKfyve knockout or inhibition alone decreased PDAC development in the KPC murine model. Mechanistically, we identify and characterize a novel relationship of synthetic lethality between PIKfyve and fatty acid synthesis. Further, we show that PIKfyve and KRAS-MAPK have a bidirectional synthetic lethality relationship: 1) PIKfyve inhibition disrupts PDAC autophagy and lysosomal function, requiring PDAC to upregulate and depend on *de novo* fatty acid synthesis through FASN and ACC1; 2) KRAS-MAPK inhibition decreases expression of FASN and ACC1 and increases PDAC utilization and reliance on autophagy; and 3) dual inhibition of PIKfyve and KRAS-MAPK drives PDAC into a metabolic crisis **(Fig. 5)**. Given the rapidly evolving landscape of mutant-KRAS^39,43^, pan-(K)RAS^68,69^, and MAPK pathway inhibitor development, this highlights the combination of PIKfyve and KRAS-MAPK inhibitors as an extremely promising and rapidly translatable therapeutic strategy for PDAC.

**Figure 5:**
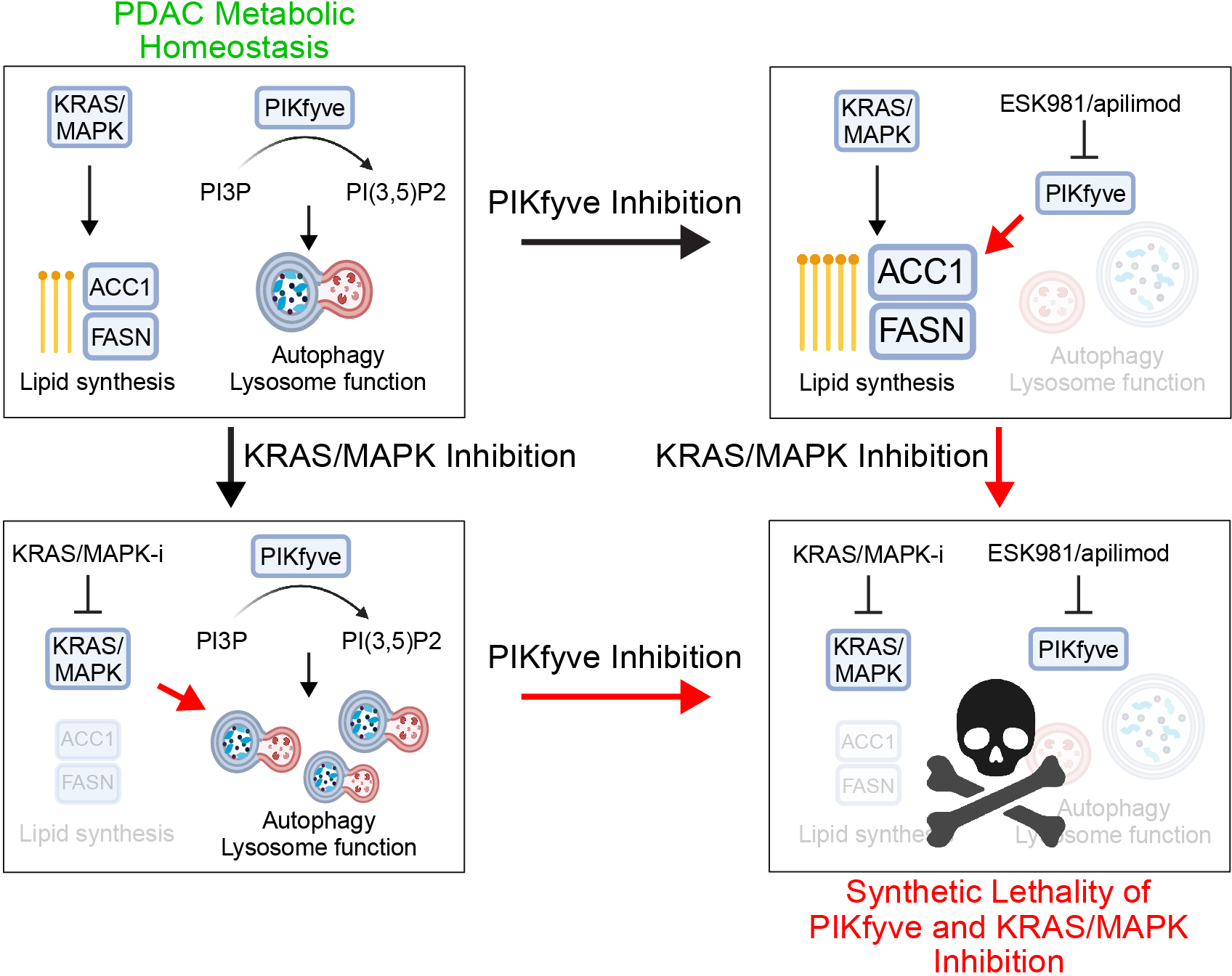
Schema depicting the effects of PIKfyve inhibition and KRAS-MAPK inhibition. (Top left) With functional PIKfyve and KRAS-MAPK signaling, PDAC is at metabolic homeostasis, able to generate lipids both through *de novo* synthesis as well as through lysosomal processes. (Top right) Upon PIKfyve inhibition, autophagy and lysosomal functions are disrupted, forcing PDAC to upregulate and depend on *de novo* fatty acid synthesis through FASN and ACC1. (Bottom left) KRAS-MAPK inhibition decreases expression of FASN and ACC1 and increases PDAC cells’ dependence on autophagy and lysosomal processes. (Bottom right) concurrent PIKfyve and KRAS-MAPK inhibition results in lethal metabolic crises in PDAC.

## Author Contributions

C.C., Y.Q., C.A.L., and A.M.C. conceived the study, designed the experiments, and composed the manuscript. Y.Q., C.A.L., and A.M.C. supervised the project. C.C. performed all *in vitro* experiments with assistance from J.P.W., Y. Zheng, L.X., D.A., and Y.B. C.C. and Y.Q. carried out all *in* vivo experiments with assistance from T.H. and Y. Zheng. P.M. carried out lipidomics experiments and analysis with assistance from R.B. Bioinformatic analyses were carried out by C.C., R.B., N.J.R., B.M., and Y. Zhang. J.H., R. Mannan, and R. Mehra carried out all histopathological evaluations and performed immunohistochemistry with assistance from C.C. and Y.Q. X.C. generated next-generation sequencing libraries and performed the sequencing. C.L., Z.W., and K.D. developed the novel PIKfyve degrader. V.S., M.P.d.M., and Y.M.S. assisted with manuscript writing and organization. M.P.d.M., Y.M.S., C.A.L., and A.M.C. provided resources and funding.

## Acknowledgements

We gratefully acknowledge Ahmet K. Korkaya, Sarah Yee, Xia Jiang, Bailey Jackson, Heng Zheng, Sydney Peters, Damien Sutton, Nupur Das, Stephanie Simko, and Andrew Delekta for technical assistance; Fengyun Su and Rui Wang for generating sequencing libraries; Somnath Mahapatra, Lisa McMurry, Amanda Miller, Christine Caldwell-Smith, Yunhui Cheng, and Shuqin Li for processing of histological samples; Jean Ching-Yi Tien for assistance with coordinating GEMM breeding colonies; Li Zhang and Anthony Andren for conducting metabolomics experiments; Lois Weisman, Daniel Klionsky, Sarah Kang, Brandon Chen, Ahmed M. Elhossiny, Megan Radyk, Samuel A. Kerk, Lin Lin, and Jae Eun Choi for experimental and analytical guidance; Agilent Technologies, Inc. for assisting with lipidomics experiments and chemometrics data analysis; Stephanie Miner for manuscript editing and preparation, and members of the Chinnaiyan and Lyssiotis laboratories and the entire Pancreatic Disease Initiative at the Rogel Cancer Center, University of Michigan, for their insightful discussions.

## Extended Data Figure Legends

**Extended Data Fig. 1: PIKfyve is essential for progression of precursor PanIN lesions to PDAC.**

A. Representative image of one additional human PDAC patient sample showing H&E (left and middle) or *PIKFYVE* RNA-ISH (right). Scalebars are 200μm (left), 20μm (middle), 20μm (right, low magnification), and 10μm (right inset, high magnification).

B. Pancreas tissue weight normalized to total body weight for *Ptf1a-Cre;Pikfyve^+/+^*, *Ptf1a-Cre*;*Pikfyve^f/+^*, and *Ptf1a-Cre*;*Pikfyve^f/f^*mice. (One-way ANOVA with Dunnett’s)

C. Representative images of H&E and insulin IHC staining from the pancreas tissue of *Ptf1a-Cre;Pikfyve^+/+^*, *Ptf1a-Cre*;*Pikfyve^f/+^*, and *Ptf1a-Cre*;*Pikfyve^f/f^* mice. Scalebar = 50μm.

D. Representative images of PIKfyve BaseScope staining from pancreas tissue of 27-week-old KC *Pikfyve^+/+^,* KC *Pikfyve^f/+^,* and KC *Pikfyve^f/f^* mice. Scalebar = 60μm for zoomed-out images; 30μm for zoomed-in images.

E. *Pikfyve* levels as determined by BaseScope of KC *Pikfyve^+/+^,* KC *Pikfyve^f/+^,* and KC *Pikfyve^f/f^* murine pancreas tissue separated by normal and lesional areas. (One way ANOVA with Dunnett’s multiple comparisons test between the indicated groups).

F. Pancreas tissue weight from 27-week-old KC *Pikfyve^+/+^,* KC *Pikfyve^f/+^,* KC *Pikfyve^f/f^*, and age-matched wild-type (WT) mice. (One-way ANOVA with Dunnett’s)

G. Pancreas tissue weight normalized to total body weight (left) and raw pancreas tissue weight (right) from 40-week old KC *Pikfyve^+/+^,* KC *Pikfyve^f/+^,* and KC *Pikfyve^f/f^*, and age-matched wild-type (WT) mice. (One-way ANOVA with Dunnett’s)

H. Percentage of pancreas occupied by normal tissue as determined by histological analyses of KC *Pikfyve^+/+^,* KC *Pikfyve^f/+^,* and KC *Pikfyve^f/f^*mice at 40 weeks of age. (One-way ANOVA with Dunnett’s)

I. Representative images of PIKfyve BaseScope staining from pancreas tissue of 25-week-old KPC *PIKfyve^+/+^*and KPC *PIKfyve^f/f^* mice. Scalebar = 20μm

J. The age at death of mice in KPC *PIKfyve^+/+^* and KPC *PIKfyve^f/f^*cohorts that were analyzed in Fig. 1O-P. (Unpaired two-tailed t-test).

**Extended Data Fig. 2: Pharmacological inhibition of PIKfyve blocks pancreatic cancer progression *in vivo*.**

A. Tumor volumes of subcutaneous CDX model derived from BxPC-3 cells in response to vehicle or ESK981 in SCID mice. Data plotted are mean tumor volumes +SEM (n=10 for each cohort). (Two-way ANOVA)

B. Individual weights (left) and images (right) of tumors from CDX model derived from BxPC-3 cells at endpoint. (Unpaired two-tailed t-test)

C. Kaplan–Meier estimates of time to tumor tripling of BxPC-3 CDX tumors after vehicle or ESK981 treatment. Statistics were performed using a Gehan-Breslow-Wilcoxon test.

D. Representative images of H&E and Ki67 IHC staining in MIA-PaCa-2 (left) and BxPC-3 (right) CDX models post vehicle or ESK981 treatment.

E. Representative image (one of three) of TUNEL staining from MDA-PaCa-2 CDX tumors after 5 days of treatment of vehicle or ESK981.

F. Tumor volumes of subcutaneous CDX model derived from T24 cells in response to vehicle or ESK981 in SCID mice. Data plotted are mean tumor volumes + SEM (n=6 for each cohort). (Two-way ANOVA)

G. Individual weights (top) and images (bottom) of tumors from CDX model derived from T24 cells at endpoint. (Unpaired two-tailed t-test)

**Extended Data Fig. 3: Genetic or pharmacological perturbation of PIKfyve disrupts autophagic flux and induces vacuolization in PDAC cells.**

A. qPCR of MIA PaCa-2 (left) or PANC-1 (right) cells upon CRISPRi-mediated knockdown of *PIKFYVE* with two independent sgRNAs (sgPIKFYVE-1 and sgPIKFYVE-2) validating target knockdown compared to control (sgNC). Data shown are technical replicates from one of three independent experiments (Unpaired two-tailed t-test).

B. Immunoblot of UM-2 primary cell-derived xenograft tumors after 5 days of treatment with either ESK981 (30 mg/kg) or vehicle as described in Figure 2K-M showing changes in LC3A/B. Vinculin was used as a loading control.

C. Tandem fluorescent reporter assay in Panc 04.03 cells showing changes in autophagic flux upon 4-hour pre-treatment with DMSO, apilimod (300 nM), ESK981 (1000 nM), or chloroquine (30 mM) and subsequent treatment of torin-1 (100 nM) or DMSO for 24 hours (one-way ANOVA with Dunnett’s). This experiment was performed independently three times with similar results.

D. Chemical structures of PIK5-12d and PIK5-33d. Red indicates protein of interest ligand; black indicates chemical linker; blue indicates VHL E3 ligase ligand.

E. Immunoblot of 7940B and MIA PaCa-2 cells treated with PIK5-33d (33d) at the indicated doses for 24 hours showing changes in p62 and LC3A/B. Vinculin was used as a loading control. These experiments were performed two times independently.

F. 20x bright-field images of MIA-PaCa-2 (top) and PANC-1 (bottom) cells upon CRISPRi-mediated knockdown of *PIKFYVE* (sgPIKFYVE-1) or control (sgNC). These images are one of three representative images.

G. 20x bright-field images of 7940B (top) MIA-PaCa-2 (bottom) cells upon treatment with PIKfyve inhibitors apilimod or ESK981 or PIK5-33d for 4 hours. These images are one of three representative images.

**Extended Data Fig. 4: Perturbation of PIKfyve decreases cell proliferation and viability *in vitro*.**

A. Confluence assay of PANC-1 cells upon CRISPRi-mediated knockdown of *PIKFYVE* (sgPIKFYVE-1 and sgPIKFYVE-2) or control (sgNC). Data shown are mean +/- SEM (n=4) from one of three independent experiments (two-way ANOVA).

B-D. Dose-response curve series of indicated PDAC cell lines treated with apilimod (B), ESK981 (C), or chloroquine (D) for 7 days using Cell-TiterGlo assays. Data presented are mean +/- SD (n=6).

E. IC_50_ values of each drug in each cell line tested.

**Extended Data Fig. 5: PIKfyve inhibition does not inhibit PDAC cell growth through disrupting iron homeostasis and mitochondrial respiration.**

A. Immunoblot of 7940B cells treated as indicated assessing changes in HIF1α, p62, and LC3A/B. Vinculin was used as a loading control.

B. Oxygen consumption rate (OCR) in 7940B (left) and Panc 04.03 (right) cells upon treatment with apilimod (100 nM), ESK981 (1000 nM), bafilomycin (BAF, 100 nM), or CQ (100 μM) for 8 hours. Automated addition of Rot/AA and 2-DG were performed at the indicated time points. Statistics were performed using one-way ANOVA with Dunnett’s multiple comparison test with DMSO as a baseline for each individual time point. Data shown represent the mean OCR +/- SEM from 5 biological replicates (One-way ANOVA with Dunnett’s, using DMSO as a reference). This experiment was performed twice independently with similar results.

C. (Left) Real-time oxygen consumption rate monitoring by Resipher on 7940B cells upon treatment with apilimod (100 nM), ESK981 (1000 nM) bafilomycin (100 nM), or chloroquine (100 μM). Data shown are mean +/-SEM from 4 biological replicates. (Right) OCR at 8 hours and 24 hours by Resipher measurements from the same experiment. (One-way ANOVA with Dunnett’s, using DMSO as a reference)

D. Dose-response curve series of 7940B (left) and PANC-1 (right) cells treated with bafilomycin, apilimod, or ESK981 in the presence or absence of ferric ammonium citrate (FAC). All conditions containing FAC were also treated with ferrostatin-1 (1 μM) to block incidental ferroptosis.

E. IC_50_ values of each treatment in cell lines with or without FAC co-treatment.

F. Confluence assay of Panc 04.03 cells undergoing treatment with deferoxamine (DFO) (100 μM), apilimod (300 nM), ESK981 (1000 nM), or bafilomycin (100 nM), without (top) or with (bottom) FAC (100 μg/mL) and ferrostatin-1 (1 μM). DFO, an iron chelator, was used as a positive control.

**Extended Data Fig. 6: Metabolic CRISPR screen nominates the *de novo* fatty acid synthesis pathway as synthetically critical upon PIKfyve inhibition.**

A. Schematic of the metabolism-focused CRISPR screen in MIA PaCa-2 cells.

B. Receiver operator characteristic (ROC) curves for the prediction of core essential genes using datasets from MIA PaCa-2 CRISPR screens.

C. qPCR showing changes in mRNA levels of *FASN* upon CRISPRi-mediated knockdown of *FASN* in MIA PaCa-2 cells. Data shown are technical triplicates from one of two independent experiments. (One-way ANOVA with Dunnett’s)

D. Confluence assays of MIA PaCa-2 cells upon *FASN* knockdown with two independent sgRNAs targeting *FASN* or control with or without treatment with PIKfyve degrader PIK5-33d (100 nM). Data shown are mean +/- SEM from 4 biological replicates. (Two-way ANOVA with Dunnett’s). This data is representative of three independent experiments. The 33d condition data was shared across both plots as they were generated in the same experiment. The DMSO condition was utilized as a control in Fig. 3I as these data were generated in the same experiment.

E. Immunoblots of PANC-1 and 7940B cells upon treatment with ND646 at indicated doses for 24 hours showing changes in labeled targets.

F. Confluence assays of MIA PaCa-2, PANC-1, and 7940B cells upon treatment with ND646 (100 nM for MIA-PaCa2, 1000 nM for PANC-1 and 7940B) and ESK981 (30 nM for MIA PaCa-2, 100 nM for PANC-1 and 7940B). Data shown are mean +/- SEM from 4 biological replicates. p-values were performed using a two-way ANOVA with Dunnett’s multiple comparisons test with the combination condition as the comparison group. These experiments were performed three independent times each. The DMSO and ND646 conditions for MIA PaCa-2 and PANC-1 are also utilized as controls in Fig. 3J and Extended Data Fig. 6G as they were generated in the same experiment. The DMSO and ND646 conditions for 7940B are also utilized as controls in Extended Data Fig. 6G as they were generated from the same experiment.

G. Confluence assays of MIA PaCa-2, PANC-1, and 7940B cells upon treatment with ND646 (100 nM for MIA-PaCa2, 1000 nM for PANC-1 and 7940B) and PIK5-33d (100 nM for MIA PaCa-2 and PANC-1, 1000 nM for 7940B). Data shown are mean +/- SEM from 4 biological replicates. p-values were determined using a two-way ANOVA with Dunnett’s multiple comparisons test with the combination condition as the comparison group. These experiments were performed three independent times each. The DMSO and ND646 conditions for MIA PaCa-2 and PANC-1 are also utilized as controls in Fig. 3J and Extended Data Fig. 6F. The DMSO and ND646 conditions for 7940B are also utilized as controls in Extended Data Fig. 6F as they were generated in the same experiment.

**Extended Data Fig. 7: PIKfyve inhibition stimulates a lipogenic transcriptional program.**

A. Scatter plot of log2 fold change in gene expression upon 8-hour treatment with apilimod (100 nM) vs DMSO (x-axis) and ESK981 (1000 nM) vs DMSO (y-axis). A linear regression was calculated with r- and p-values displayed.

B. GSEA plots of cholesterol homeostasis and fatty acid metabolism using the fold change rank-ordered gene signature from the 7940B cells treated with apilimod or ESK981 for 8 hours.

C. Volcano plot using RNA-seq analysis on 7940B cells treated with ESK981 (1000 nM) for 8 hours highlighting SREBP-1 target genes.

**Extended Data Fig. 8: Inhibition of PIKfyve induces lipid synthesis and accumulation of sphingolipids.**

A. Principal component analysis (PCA) of targeted metabolomics experiment on 7940B cells treated with DMSO, apilimod (100 nM), or ESK981 (1000 nM) for 8 hours.

B. Citrate levels as detected by LC-MS-based metabolomics on 7940B cells treated with apilimod (100 nM) or ESK981 (1000 nM) for either 3 or 8 hours, as indicated. Data shown are biological triplicates (one-way ANOVA with Dunnett’s).

C. Heatmap of significantly changed metabolites as determined by unpaired two-tailed t-test (apilimod vs DMSO or ESK981 vs DMSO, *p < 0.05* in at least one of the two comparisons). Biological triplicates are displayed.

D. PCA of targeted lipidomics experiment on 7940B cells treated with DMSO, apilimod (100 nM), or ESK981 (1000 nM) for 24 hours.

E. Volcano plot showing differentially abundant lipid species in 7940B cells upon treatment with ESK981 (1000 nM) for 24 hours. Highlighted in red are upregulated sphingolipid species. p-value was determined by unpaired two-tailed t-test. Highlighted in blue are downregulated sphingolipid species.

**Extended Data Fig. 9: KRAS-MAPK perturbation downregulates lipogenic transcriptional program and induces autophagic flux in direct opposition to PIKfyve inhibition’s effects.**

A. qPCR of iKRAS (doxycycline-inducible KRAS^G12D^) 9805 cells showing changes in RNA levels of labeled genes upon presence or absence of doxycycline (Dox) for 48 hours. Data plotted are technical triplicates. This experiment was performed three independent times. (Unpaired two-tailed t-test)

B. qPCR of 7940B and MIA PaCa-2 cells treated with KRAS^G12D^ inhibitor MRTX1133 (100 nM), KRAS^G12C^ inhibitor AMG510 (100 nM for MIA PaCa-2), or MEK inhibitor trametinib (10 nM for MIA PaCa-2, 30 nM for 7940B) for 8 hours. Data plotted are technical triplicates from one of three independent experiments. (One-way ANOVA with Dunnett’s)

C. Counts per million (CPM) from RNA-seq analysis on AsPC1 cells treated with MRTX1133 (100 nM) for 24 hours. Data plotted are biological triplicates from publicly available RNA-seq data^43^. (Unpaired two-tailed t-test)

D. CPM from RNA-seq analysis on AsPC1 cell-derived xenograft model. Mice were dosed with 30 mg/kg of MRTX1133 6 hours prior to tumor collection. Data plotted are taken from independent tumors from publicly available RNA-seq data^43^. (Unpaired two-tailed t-test)

E. Tandem fluorescent reporter assay in 7940B (left) or Panc 04.03 (right) cells showing changes in autophagic flux upon 24-hour treatment with labeled doses of MRTX1133. Data shown are 4 biological replicates from one of three independent experiments. (One-way ANOVA with Dunnett’s)

F. Tandem fluorescent reporter assay in Panc 04.03 cells showing changes in autophagic flux upon 4-hour pretreatment with apilimod (300 nM), ESK981 (1000 nM), or chloroquine (CQ, 30 mM) followed by 24-hour treatment with MRTX1133 (300 nM, left) or trametinib (25 nM). Data shown are 4 biological replicates from one of three independent experiments. (One-way ANOVA with Dunnett’s)

**Extended Data Fig. 10: Dual inhibition of PIKfyve and KRAS-MAPK results in synergistic cell growth suppression *in vitro*.**

A. 3D synergy plots in 7940B cells treated with ESK981 and trametinib (left), apilimod and MRTX1133 (middle), and ESK981 and MRTX1133 (right). Red peaks in the 3D plots indicate synergism, and the overall average synergy score is listed above.

B. Corresponding heatmaps for Extended Data Fig. 10A of cell viability decreases upon individual or combinatorial treatments from five biological replicates each. Relative viability from each plate were normalized to the viability of 10 untreated wells in each respective plate.

C. Confluence assays of 7940B cells treated with DMSO, trametinib (20 nM), apilimod (50 nM), or both trametinib and apilimod. Data shown are mean +/- SEM from 4 biological replicates. Statistics were performed using a two-way ANOVA with Dunnett’s multiple comparisons test.

**Extended Data Fig. 11: Combination therapy of trametinib and ESK981 is well tolerated in immune competent mice and results in elimination of tumor burden in a syngeneic orthotopic model.**

A. Relative body weight (compared to day 1) of mice harboring 7940B orthotopic tumors undergoing indicated treatment. Data displayed are average body weight of each cohort +/- SEM from all mice utilized in the study (vehicle = 9, tram= 9, ESK= 9, ESK+tram = 8).

B. Raw pancreas + tumor weights collected at endpoint of 7940B syngeneic orthotopic model. (One-way ANOVA with Tukey’s)

C. Barplot of tumor presence or absence based on histological evidence.

## Supplementary Tables

**Supplementary Table 1: sgRNA counts for metabolism-focused CRISPR screen.**

**Supplementary Table 2: Antibodies, primers, sgRNAs, and compounds used in this study.**

**Supplementary Table 3: Processed data in transcripts per million (TPM) for RNA-seq experiment.**

**Supplementary Table 4: Source data for metabolomics experiment.**

**Supplementary Table 5: Source data for lipidomics experiment.**

## Methods

### Mouse strains

*Ptf1a-Cre, Ptf1a-Cre; lsl-Kras^G12D^* (KC), and *Ptf1a-Cre; lsl-Kras^G12D^; p53^R172H/+^* (KPC) mice were previously described^70,71^. Conditionally floxed *Pikfyve* (*Pikfyve^f/f^*) mice were purchased from Jackson labs. PCR genotyping for *Ptfia-Cre, Kras^G12D^*, *p53^R172H/+^*, and *Pikfyve^f/f^ alleles*, from DNA isolated from mouse tails, was performed using standard methodology. Littermate controls were systematically used in all experiments, and the sex ratios for each cohort were balanced. All animals were housed in a pathogen-free environment, and all procedures were performed in accordance with requirements of the University of Michigan Institutional Animal Care & Use Committee (IACUC).

### Cell lines, antibodies, and compounds

PANC-1, MIA PaCa-2, Panc 04.03, SW1990, Panc 10.05, and HPAF-II were originally obtained from the American Type Culture Collection (ATCC). 7940B was generously provided by Greggory Beatty, M.D., Ph.D. at Perlman School of Medicine at the University of Pennsylvania. The iKRAS 9805 cell line was previously described^72^. The UM PDAC primary cell lines (UM2, UM19) were obtained from surgically resected samples and established through murine xenograft^73^. KPC-1344 was derived from a KPC mouse in-house by dissociating tumors manually with a sterile blade and then treating them with 1 mg/mL collagenase II (ThermoFisher Scientific cat. no. 17101-015) and 1 mg/mL DNase (Sigma-Aldrich, cat. no. 10104159001) for 30 minutes with shaking at 37°C. The cells were then strained using a MACS SmartStrainer (30μM) (Miltenyi Biotec cat no. 130-110-915) and rinsed with PBS prior to culturing. All cells were grown in Gibco DMEM + 10% FBS (ThermoFisher). All cell lines were genotyped to confirm their identity by Eurofins Genomics and tested biweekly for mycoplasma contamination. Sources of all antibodies and compounds are described in **Supplementary Table 2**.

### Histopathologic analyses

The study pathologists conducted a detailed histopathological evaluation of murine pancreatic tissues on 4 μm thick H&E-stained formalin fixed paraffin embedded (FFPE) sections. The examination involved checking all harvested pancreas samples for the percentage prevalence of normal pancreas, pancreatic intraepithelial neoplasia (PanIN)- either high and low grade, and lesions with atypia or frank evidence of pancreatic ductal adenocarcinoma. The samples were then classified under these three categories, and the results were tabulated. Finally, the two pathologists reached a consensus to determine the final percentage prevalence.

### *PIKFYVE* RNAScope

RNA-ISH was performed using the RNAscope 2.5 HD Brown kit (Advanced Cell Diagnostics/ACD, Newark, CA) and target probe against *PIKFYVE* (Cat No. 1326631 Hs- *PIKFYVE*) according to the manufacturer’s instructions. RNA quality was evaluated in each case utilizing a positive control probe against human housekeeping gene Peptidylprolyl Isomerase B (PPIB) (Cat No. 313901). Assay background was monitored using a negative control probe against bacillus bacterial gene DapB (Cat No. 310043). Stained slides were evaluated under a light microscope at low- and high-power magnification for RNA-ISH signals in the cancer cells and normal pancreas by multiple study investigators (R. Mannan, and J. Hu). The expression level was evaluated according to the RNAscope scoring criteria: score 0 = no staining or <1 dot per 10 cells; score 1 = 1–3 dots per cell, score 2 = 4–9 dots per cell, and no or very few dot clusters; score 3 = 10-15 dots per cell and <10% dots in clusters; score 4 = >15 dots per cell and > 10% dots in clusters. The RNA-ISH score was calculated for each examined tissue section as the sum of the percentage of cells with score 0-4 [(A% × 0) + (B% × 1) + (C% × 2) + (D % × 3) + (E% × 4), A + B + C + D + E = 100], using previously published scoring criteria^74^.

### *Pikfyve* BaseScope

The BaseScope™ VS Reagent Kit (Cat. No. 323700; Advanced Cell Diagnostics, Newark, CA), which is used to identify short targets and splice variants, was employed to demonstrate *Pikfyve* on whole mouse pancreatic tissues. The reagent kit was used with the Discovery Ultra automated IHC/ISH slide staining systems by Ventana Medical Systems on a validated protocol utilizing BaseScope™ VS Detection Reagents (Cat. No. 323710), RNAscope Universal VS Sample Preparation Reagents v2 (Cat. No. PN323740), and RNAscope VS Accessory Kit (320630). BaseScope™ VS Probe - BA-Mm-Pikfyve-E6-3zz-st-C1, Mus musculus phosphoinositide kinase FYVE type zinc finger containing (Pikfyve) transcript variant 2 mRNA targeting exon 6 complimentary to the target mRNA was employed (Cat. No. 1300097-C1; accession # NM_011086.2, nucleotides 633-771) for the assay as test probe. BaseScope™ VS Positive Control Probe -Mm-PPIB-3ZZ - Mus musculus peptidylprolyl isomerase B (Ppib)mRNA (Cat. No701079) and BaseScope™ VS Negative Control Probe-DapB-3ZZ (Cat No. 701019) were used as positive and negative controls, respectively.

All slides were examined for positive signals in lesions and background benign pancreas by 2 study pathologists (R. Mannan and J. Hu). The RNA *in situ* hybridization signal was identified as red, punctate dots, and the expression level was scored as follows: 0=no staining or <1 dot per 10 cell (at 40X magnification), 1= 1 dot per cell (visible at 20/40X), 2= 2-3 dots per cell, 3=4-10 per cell (<10% in dot clusters) visible at 20X, and 4=>10 dots per cell (>10% in dot clusters) visible at 20X. A cumulative RNA ISH product score (BaseScope score) was calculated for each evaluable tissue core as the sum of the individual products of the expression level (0 to 4) and percentage of cells [0 to 100; ie, (A%×0)+(B%×1)+(C%×2)+(D%×3)+(E%×4); total range=0 to 400]

### Immunohistochemistry

Immunohistochemistry was performed on formalin-fixed paraffin-embedded 4 μm sections of mouse or xenograft tissues. Slides were deparaffinized in xylene, followed by serial hydration steps in ethanol (100%, 95%, 70%) and water for 4 minutes each. Antigen retrieval was performed by boiling slides in citrate buffer (pH 6). Endogenous tissue peroxidase activity was blocked by 3% H2O2 for 1 hour. Slides were blocked in 10% goat serum for 1 hour. The slides were then incubated in the primary antibodies. The specifics of the antibodies used are listed in **Supplementary Table 2**. Visualization of staining was done per the manufacturer’s protocol (Vector Laboratories, cat. no. SK-4100). Following DAB staining, slides were dehydrated in ethanol (70%, 95%, 100%, 6 minutes each), xylene (15 minutes), and mounted using EcoMount (Thermo Fisher, cat. no. EM897L).

Following IHC staining, quantification was carried out using Fiji (Imagej)^75^ (Fig. 4L). Images were first subjected to color deconvolution using the H DAB vector. Subsequently, a manual threshold was set based on the uniform signal intensity of the DAB signal, serving as a cut-off for all images. The ratio of brown signal to total signal was calculated as the CK19% positive area displayed on the figure. Regions outside the pancreas, such as the spleen, were excluded from the analysis.

### *In vivo* tumor studies

All animal experiments were conducted in accordance with the Office of Laboratory Animal Welfare and approved by the University of Michigan IACUC.

#### Subcutaneous tumor studies

For xenograft studies, 6-8-week-old CB17 severe combined immunodeficiency (SCID) mice obtained from the University of Michigan breeding colony were used. For syngeneic studies, 6-8-week-old C57BL6 mice obtained from Jackson Laboratories were used. Subcutaneous tumors were established at both sides of the dorsal flank of the mice by injecting 1×10^6^ cells in 100 μL of 50:50 Matrigel and serum-free media. Tumors were measured 2-3 times per week using digital calipers following the formula (π/6) (L× W2), where L = length and W = width of the tumor. At the end of the studies, mice were sacrificed, and tumors extracted and weighed.

#### Pancreatic orthotopic tumor study

The 7940B orthotopic model was established according to previously described protocols^15^. Briefly, 50,000 cells were implanted directly into the pancreas of C57BL/6J mice (Jackson Laboratories). Tumors were established for 11 days prior to treatment with the indicated conditions. Mice were sacrificed at 3 weeks of treatment, and tumors were weighed and preserved for further analyses.

### *In vivo* apoptosis evaluation using TUNEL staining

Terminal dUTP Nick End Labeling (TUNEL) staining was performed with an *In Situ* Cell Death Detection Kit (TMR Red #12156792910; Roche Applied Science) following the manufacturer’s instructions. Briefly, fixed sections were deparaffined, rehydrated, and subsequently permeabilized using proteinase K. The labelling reaction was performed at 37°C for 1 hour by addition of the reaction buffer containing enzymes. Images were acquired using a Zeiss Axiolmager M1 microscope. Quantification was performed using Fiji (ImageJ)^75^ (Fig. 2K). Signal from TUNEL and from DAPI were quantified independently using the same manual threshold for all samples. %TUNEL positive scores were calculated as a percentage of TUNEL signal divided by DAPI signal.

### Immunoblots

Cell lysates were prepared in RIPA buffer (ThermoFisher Scientific) supplemented with Halt™ Protease and Phosphatase Inhibitor Cocktail (ThermoFisher Scientific). Total protein was measured by DC™ Protein Assay Kit II (BIO-RAD). An equal amount of protein was resolved in NuPAGE™ 3 to 8%, Tris-Acetate Protein Gel (ThermoFisher Scientific) or NuPAGE™ 4 to 12%, Bis-Tris Protein Gel (ThermoFisher Scientific), blocked with 5% nonfat dry milk in TBS-T and blotted with primary antibodies overnight. Following incubation with HRP-conjugated secondary antibodies, membranes were imaged on an Odyssey Fc Imager (LiCOR Biosciences). For immunoblot experiments involving multiple targets overlapping in size, sample lysates were prepared in bulk and loaded on multiple gels as needed. One representative loading control for each experiment was displayed on the figures.

### Cellular Thermal Shift Assay (CETSA)

CETSA was performed according to previously described protocols^76^. Briefly, 7940B cells were seeded overnight and subsequently treated with DMSO, ESK981 (1000 nM), or apilimod (1000 nM) for 2 hours. Cells were then harvested and made into single-cell suspensions of 1×10^6^ cells each in 50 μL of PBS containing protease inhibitors. The suspensions were then subjected to heating and cooling cycles (two cycles of 3-minute heating followed by 3-minute cooling at room temperature) using a thermal cycler. Cells were then lysed with three cycles of freeze-thawing in liquid nitrogen. Lysates were then centrifuged at 12,000 x g for 10 minutes, and the soluble fraction was isolated, denatured, and resolved on a NuPAGE™ 4 to 12%, Bis-Tris Protein Gel (ThermoFisher Scientific), blocked with 5% nonfat dry milk in TBS-T and blotted with primary antibodies overnight. Following incubation with HRP-conjugated secondary antibodies, membranes were imaged on an Odyssey Fc Imager (LiCOR Biosciences).

### Cell viability assays and synergy assays

Cells were plated into 96-well plates and incubated overnight at 37°C in 5% CO_2_. The following day, a serial dilution of the indicated compounds was prepared in culture medium and added to the plate. The cells were then further incubated for 5 days (experiments involving MRTX1133 or trametinib) or 7 days (all other experiments). Subsequently, the CellTiter-Glo assay (Promega), was then performed according to the manufacturer’s instructions. The luminescence signal was acquired using an Infinite M1000 Pro plate reader (Tecan), and the data were analyzed using GraphPad Prism 10 (GraphPad Software Inc.).

To determine the synergism of two different compounds using viability assays, cells were treated with the indicated combinations of the drugs for 5 days prior to performing the CellTiter-Glo assay as described above. These experiments were performed with 5 biological replicates each with 10 wells of untreated internal controls for each plate used in each experiment which were used for normalization between plates. The data were then expressed as percent inhibition relative to baseline, and the presence of synergy was determined by the Bliss method using the SynergyFinder+ web application^77^.

### Autophagic flux probe generation and assay

Generation of the autophagic flux probe in 7940B, Panc 04.03, and iKRAS was done according to the original author’s instructions^60^. Briefly, cells were infected with pMRX-IP-GFP-LC3-RFP-LC3ΔG, which was a gift from Noboru Mizushima (Addgene #84572). Following puromycin selection, single-cell clones were expanded and genotyped to ensure the absence of homologous recombination between two LC3 fragments during retrovirus integration.

15,000 cells were seeded in 96-well plates. After overnight incubation, cells were treated with the indicated compounds for 24 hours. For assays assessing co-treatment of autophagy inhibitors (i.e., apilimod, ESK981, chloroquine) with autophagy inducers (torin-1, trametinib, MRTX1133), the autophagy inhibitor was added 4 hours prior to the inducer. For assays using iKRAS, cells were seeded with or without doxycycline, as indicated, and then treated with compounds in a similar fashion. Fluorescent signals were detected using the Infinite M1000 Pro plate reader (Tecan). Autophagy index was calculated by dividing the RFP signal by GFP signal from each well, followed by normalization of all RFP/GFP ratios by the average RFP/GFP ratio of the DMSO condition.

### Confluence-based proliferation assays (Incucyte)

Cells were seeded in a clear 96-well plate overnight prior to treatment. Upon treatment with indicated compounds, plates were incubated in an Incucyte S3 2022 Rev1 (Sartorious), with 10x images taken every 4 hours, and confluence was analyzed to assess for proliferation.

### Oxygen consumption assays

Oxygen consumption rates were determined using the Seahorse XF Glycolytic Rate Assay (Agilent) according to the manufacturer’s protocol. Briefly, 15,000 (7940B) or 25,000 (Panc 04.03) cells were seeded in an Agilent XF96 Cell Culture Microplate 16 hours prior to treatment. Cells were treated with AP, ESK, CQ, or BAF as indicated for 8 hours. Immediately prior to the assay, cells were washed and then incubated in XF DMEM medium (pH 7.4, Agilent) with 1 mM pyruvate, 2 mM glutamine, and 10 mM glucose. The assay was conducted on an XF96 Extracellular Flux Analyzer (Agilent), and the OCR was calculated using Wave (version 2.6, Agilent). OCR was normalized to cell number with the CyQUANT NF Cell Proliferaiton Assay (Invitrogen) according to the manufacturer’s instructions.

Real-time monitoring of basal oxygen consumption rate was performed using a Resipher (Lucid Scientific). 15,000 7940B cells were seeded in 50 μl of medium in a clear 96-well plate 16 hours prior to treatment. Immediately following treatment with an additional 50 μl of medium (for a total of 100 μl), OCR monitoring was started by placing the Resipher device on the cells, which was incubated at 37°C and 5% CO_2_ for 24 hours.

### Metabolic CRISPR screen

The Human CRISPR Metabolic Gene Knockout library was a gift from David Sabatini (Addgene #110066)^78^. To achieve at least 1000-fold coverage of the library while culturing, 75 x 10^6^ MIA PaCa-2 cells were seeded at a density of 5 x 10^5^ cells/mL in 6-well plates containing 2 mL of DMEM, 8 mg/mL polybrene, and the CRISPR screen library virus. Spin infection was caried out by centrifugation at 1200 g for 45 minutes at 37°C. After 24-hour incubation, the media was replaced with fresh DMEM. After a subsequent 24-hour incubation, cells were transferred to T-150 flasks (at a density of 3 wells into 1 T150 flask) containing 20 mL of DMEM containing puromycin at 2 mg/mL. After 3 days of selection, cells were seeded into sixteen total T-150 flasks at a density of 5 x 10^6^ cells/flask in 20 mL of DMEM containing either DMSO or 200 nM of apilimod. Cells were passaged every 3-4 days and re-seeded back to the original cell density. After 14 days, a pool of 15 million cells from each condition were harvested for genomic DNA (gDNA) isolation using the DNeasy blood and tissue kit (Qiagen) according to the manufacturer’s protocol.

For each condition, sgRNA was amplified from 50 mg gDNA using Herculase II Fusion DNA Polymerase (Agilent Technologies), column purified using Select-a-Size DNA Clean & Concentrator kit (Zymo Research), and then gel-purified using 6% Novex TBE gel (Thermo), followed by isolation from the gels with Gel Breaker Tubes and Gel Filters (BioChain). The resulting PCR products then underwent end-repair and A-tail addition followed by New England Biolabs (NEB) adapter ligation. The final library was prepared by enriching adapter-ligated DNA fragments using 2x KAPA HiFi HotStart mix and NEB dual code barcode following the manufacturer’s protocol. The libraries were then sequenced on an Illumina NovaSeq 6000 (paired-end 2 x 151 nucleotide read length).

Reads were trimmed to the bare sgRNA sequence using cutadapt 4.1^79^. Paired-end mates were trimmed separately using a sequence 5’-adjacent to the sgRNA position within the vector (TATATCTTGTGGAAAGGACGAAACACCG), requiring a minimum match of 18 bases to the sequence and followed by truncation to 20 bases (relevant cutadapt command parameters: *-m 18 -O 18 -l 20 --discard-untrimmed*). Trimmed reads were then combined and aligned using bowtie2 2.4.5^80^ to a reference built from each sgRNA in the library flanked by vector sequences (5’ GTTATCAACTTGAAAAAGTGGCACCG and 3’ CTAGATCTTGAGACAAATGGC). The bowtie2 parameter *--norc* was used to prevent reverse compliment alignment. Counting was then performed using MAGeCK 0.5.9.5^81^. See **SupplementaryTable 1** for a summary of read counts.

sgRNAs with less than 100 counts in the initial dataset were removed from downstream analysis. Genes targeted by fewer than 6 distinct sgRNAs following this filtering were likewise removed. Downstream analyses, including calculation of sgRNA depletion/enrichment scores, gene depletion/enrichment scores, and selective dependency, were done according to previously described methods^82^. Briefly, normalized sgRNA abundances were calculated by adding a pseudocount of one and then normalized to the total counts of each sample. The sgRNA enrichment/depletion scores were calculated as log2 fold change between the final and initial populations, and the gene scores were calculated as the average log2 fold change of the sgRNAs targeting that gene. To calculate selective essentiality scores, we first scaled gene scores using the medians of nontargeting sgRNAs and sgRNAs targeting core essential genes as references (0 and -1, respectively). Selective essential genes were then identified by taking the Z-scored difference between the scaled apilimod and DMSO gene scores. Plots were generated using ggplot2 (version 3.4.4).

### RNA isolation and quantitative real-time PCR (qPCR)

Total RNA was isolated from cells using the miRNeasy kit (Qiagen), and cDNA was synthesized from 1000 ng of total RNA using Maxima First Strand cDNA Synthesis Kit for RT-qPCR (Thermo Fisher Scientific). Quantitative real-time PCR was performed in triplicates using standard SYBR green reagents and protocols on a QuantStudio 5 Real-Time PCR system (Applied Biosystems). The target mRNA expression was quantified using the ΔΔCt method and normalized to *ACTB* (human) or *Actb* (murine) expression. All primers were synthesized by Integrated DNA Technologies (Coralville). Primer sequences are listed in **Supplementary Table 2**.

### RNA-seq and analysis

RNA-seq libraries were prepared using 800 ng of total RNA. Ribosomal RNA was removed by enzymatic digestion of the specific probe-bound duplex rRNA, and then fragmented to around 200-300 bp with heat in fragmentation buffer (KAPA RNA Hyper+RiboErase HMR, Roche). Double-stranded cDNA was then synthesized by reverse transcription and underwent end-repair and ligation using New England Biolabs (NEB) adapters. Final library preparation was then prepared by amplification with 2x KAPA HiFi HotStart mix and NEB dual barcode. Library quality was measured on an Agilent 2100 Bioanalyzer (DNA 1000 chip) for concentration and product size. Paired-end libraries were sequenced with the Illumina NovaSeq 6000, (paired-end 2 × 151 nucleotide read length) with sequence coverage to 30-40 million paired reads. Reads were demultiplexed using Illumina’s bcl2fastq conversion software v2.20. Transcripts were quantified by the alignment-free approach kallisto^83^ using index generated from mouse reference genome (mm10) and then summed to obtain gene level counts. Raw Transcripts Per Million values for each gene can be found in **Supplementary Table 3**. Differential analysis was performed using limma-voom^84,85^ after TMM-normalization^86^ of gene level counts with calcNormFactors of edgeR^87^. Genes with mean Transcripts Per Million (TPM) less than 1 in both control and treatment groups were considered as lowly expressed genes and excluded for differential analysis. Enrichment of Hallmark and Reactome gene sets downloaded from MSigDB^88^ were examined with fgsea^89^ using genes ranked by logFC estimated from limma as input.

### Generation of CRISPRi-mediated knockdown cell lines

sgRNA sequences used were taken from previously validated Perturb-seq library^90^. The sgRNAs were cloned into the backbone, pLV hU6-sgRNA hUbC-dCas9-KRAB-T2a-Puro (Addgene plasmid # 71236; http://n2t.net/addgene:71236; RRID: Addgene_71236)^91^ using the Golden Gate reaction. The generated plasmids were then expanded, verified by Sanger sequencing, and packaged into lentiviruses by the University of Michigan Vector Core. Cells were seeded, infected with viruses along with polybrene (10 mg/mL), and then selected with puromycin (2 μg/mL for MIA PaCa-2, 5 μg/mL for PANC-1) prior to further analysis. Given the notable impact of *PIKFYVE* and *FASN* knockdown on PDAC cells, new CRISPRi knockdown cell lines were generated prior to each experiment. The sgRNA sequences are listed in **Supplementary Table 2**.

### ESK981, trametinib, and MRTX1133 formulation for *in vivo* studies

ESK981 was added to ORA-PLUS and sonicated until completely dissolved. Trametinib was added to corn oil and sonicated until completely dissolved. Aliquots were frozen at -20°C to prevent freeze-thaw cycles. MRTX1133 was added to 10% Captisol in 50mM Citrate (pH = 5.0) and sonicated until completely dissolved as previously described^43^. Dissolved MRTX1133 was kept at 4°C hidden from light for a maximum for 5 days. ESK981 and trametinib were delivered by oral gavage. MRTX1133 was delivered by intraperitoneal (IP) injection.

### Targeted metabolomics

Polar metabolites from samples treated in biological triplicates were extracted using 80% v/v methanol/water and normalized using protein quantification from an additional sample from each condition. Equal estimated amounts of metabolites were dried using a SpeedVac vacuum concentrator, reconstituted in 50% v/v methanol in water, and analyzed by LC-MS as previously described^92^. Data were analyzed as previously described^92^ with the Agilent MassHunter Workstation Quantitative Analysis for QQQ version 10.1 Build 10.1.733.0. However, metabolite abundance levels were not divided by the median levels across the samples. No post-detection normalization was performed to avoid assuming linearity of signal. Raw values of each metabolite measured are provided in **Supplementary Table 4**. Heatmaps were generated using the Morpheus Matrix Visualization and analysis tool (https://software.broadinstitute.org/morpheus).

### Targeted lipidomics

Sample preparation: Samples for lipidomics analyses were prepared according to the automatic dual-metabolite/lipid sample preparation workflow described in the Agilent application note 5994-5065EN. Briefly, 1 million cells were washed in PBS and lysed with 1:1 trifluoroethanol/water at room temperature. Lysates were transferred to microcentrifuge tubes, incubated for 10 minutes, and centrifuged at 250 x g for 30 seconds. Samples were dried with a vacuum concentrator and resuspended in 1:1 trifluoroethanol/water. After transferring the samples to a 96-well plate, lipids were selectively isolated on a Bravo automated liquid handler platform (Agilent) operated by a VWorks automation protocol as described (5994-5065EN).

LC-MS/MS analysis: Samples were analyzed on an Agilent 1290 Infinity II Bio LC ultra-high performed liquid chromatography (UPLC) system with the Agilent Standardized Omics LC configuration, consisting of a high-pressure binary pump, multicolumn thermostat, and a temperature controlled multisampler. Samples were injected in randomized order on an Agilent 6495C triple quadrupole mass spectrometer equipped with an Agilent Jet Stream Dual ESI ion source. Samples were analyzed with the reverse phase LC-MS/MS method reported in the Agilent application note 5994-3747EN. After acquisition, datasets were processed with MassHunter Quantitative Analysis 12.0 software and subsequently imported into Mass Profiler Professional (MPP) for chemometric analysis. No post-detection normalization was performed to avoid assuming linearity of signal. Raw values of each lipid measured are provided in **Supplementary Table 5**.

Changes in lipid class abundance in 7940B cells upon treatment with apilimod (100 nM) or ESK981 (1000 nM) relative to treatment with DMSO were estimated from linear mixed models with random intercepts to adjust for the baseline differences across the lipid classes. A separate model for each treatment (apilimod or ESK981) comparison against DMSO is built using the R package lme4 (version 1.1-35.1)^93^.

### Statistics and reproducibility

No data were excluded from the analyses. No statistical methods were used to predetermine sample sizes. For all *in vivo* experiments, animals were randomly assigned into treatment cohorts. Tumor measurements were performed by digital caliper in a blinded manner. For all *in vitro* experiments, cells were seeded from the same pool, and, thus, there was no requirement for randomization. All samples were analyzed equally and simultaneously to eliminate bias. All error bars indicate +/-SD unless otherwise indicated. All box-and-whisker plots display the entire range of values and display all data points. All statistics comparing two groups were performed using unpaired two-tailed t-tests unless otherwise indicated. All statistics comparing more than two groups were done using an ANOVA with Dunnett’s multiple comparison test, using the indicated group as a control, unless otherwise indicated. All statistics comparing two curves were performed using a two-way ANOVA. All statistics comparing four curves were performed using a two-way ANOVA with Dunnett’s multiple comparison test, using the indicated curve as a control. GraphPad Prism software (version 10) and R v.4.3.2 were used for statistical calculations. Specific R packages utilized for individual analyses were included in their specific Methods section.

### Data and materials availability

All raw data will be provided before publication as part of sources and supplementary data files. All materials are available from the authors upon reasonable request. All raw next-generation sequencing, such as DNA sequencing for the CRISPR screen or RNA-seq, have been deposited in the Gene Expression Omnibus (GEO) repository at NCBI with the accession number GSE255378. Processed sequencing data, such as sgRNA counts and RNA-seq CPM, will be included as part of sources or supplementary data files. Raw data for metabolomics and lipidomics experiments are included as Supplementary Tables (4 and 5, respectively).

### Code availability

No custom codes were developed for this study.

### Statement on use of human samples

Patient tissues from biopsies of pancreatic tumors were acquired from the University of Michigan (U-M) pathology archives. These tissues were utilized for RNA Scope (RNA-ISH) experiments to assess for *PIKFYVE* expression in tumor or adjacent normal pancreatic cells. Use of clinical formalin-fixed paraffin embedded specimens from the archives was approved by the U-M Institutional Review Board and does not require patient consent.

