## Extended Data Figures 1 - 11 for "Targeting PIKfyve-driven lipid homeostasis as a metabolic vulnerability in pancreatic cancer"

Extended Data Fig. 1

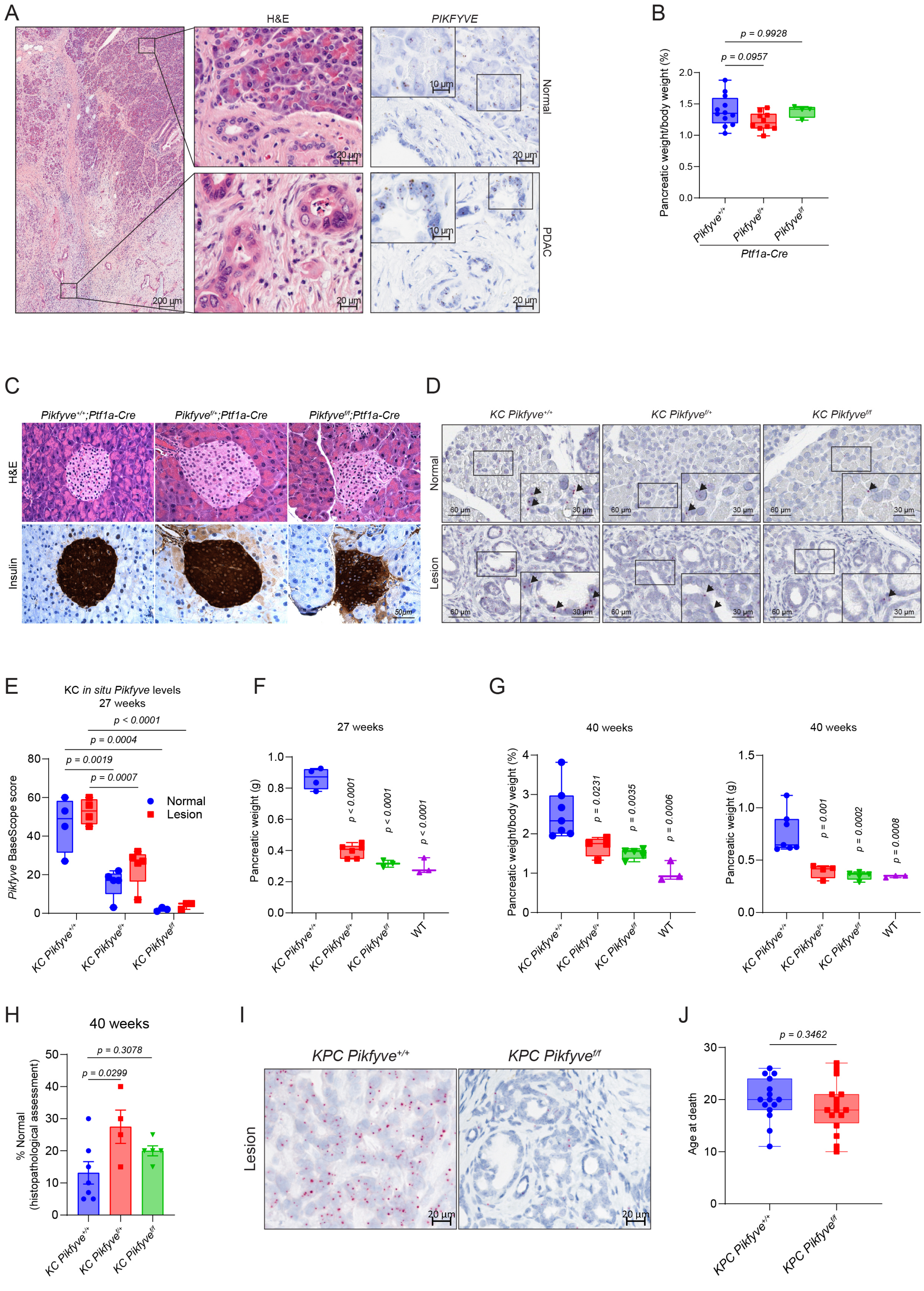

Extended Data Fig. 2

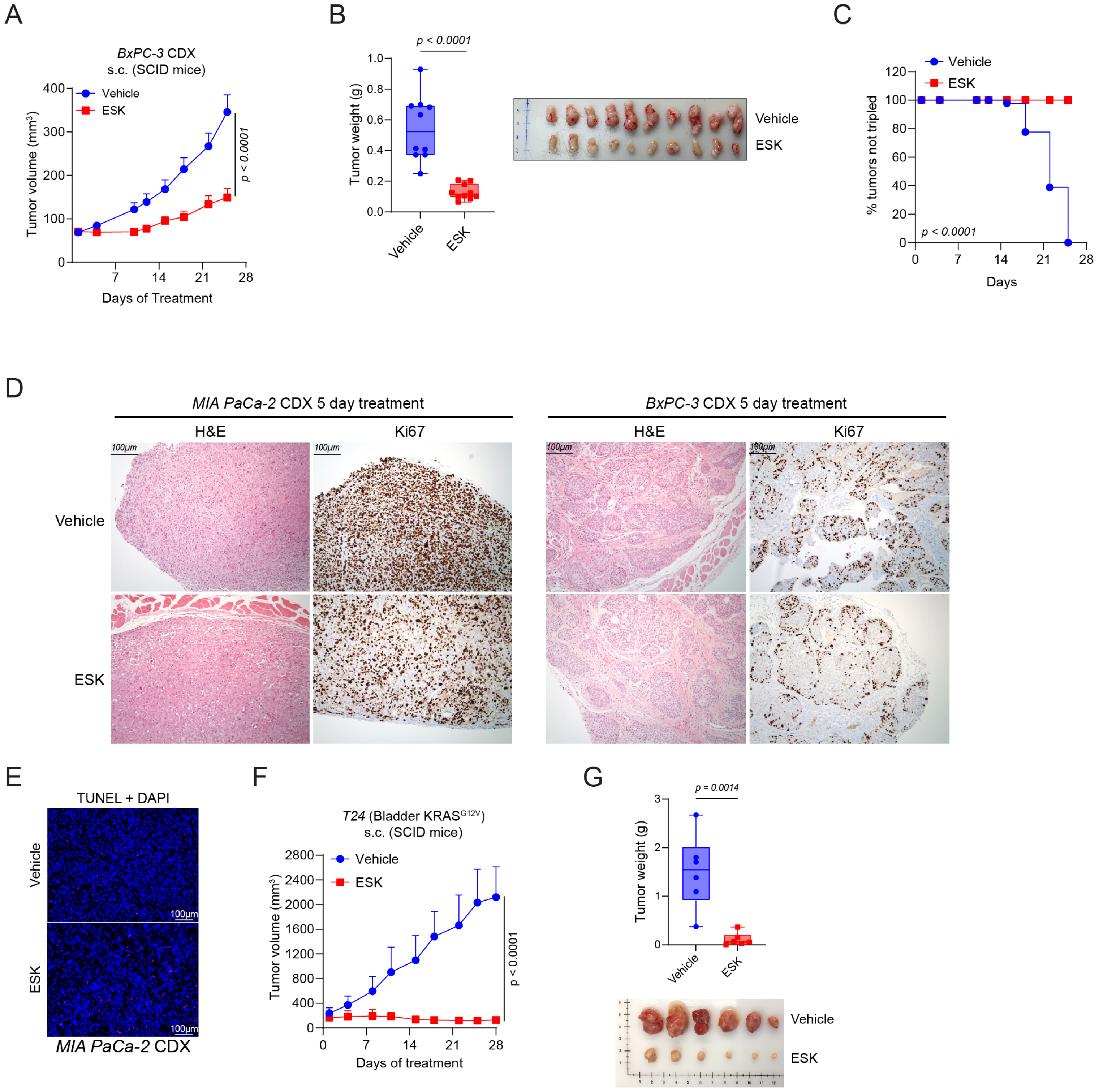

Extended Data Fig. 3

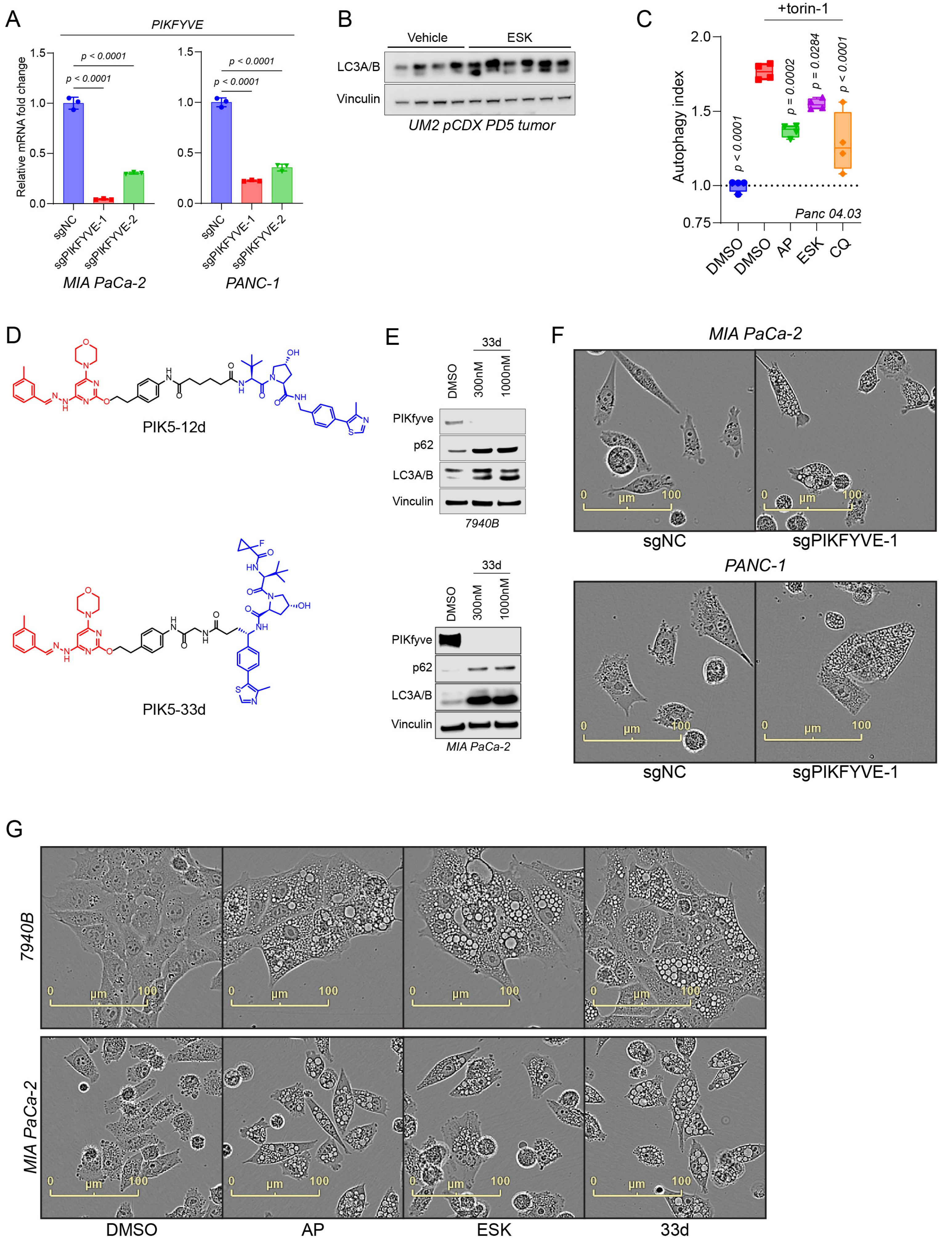

Extended Data Fig. 4

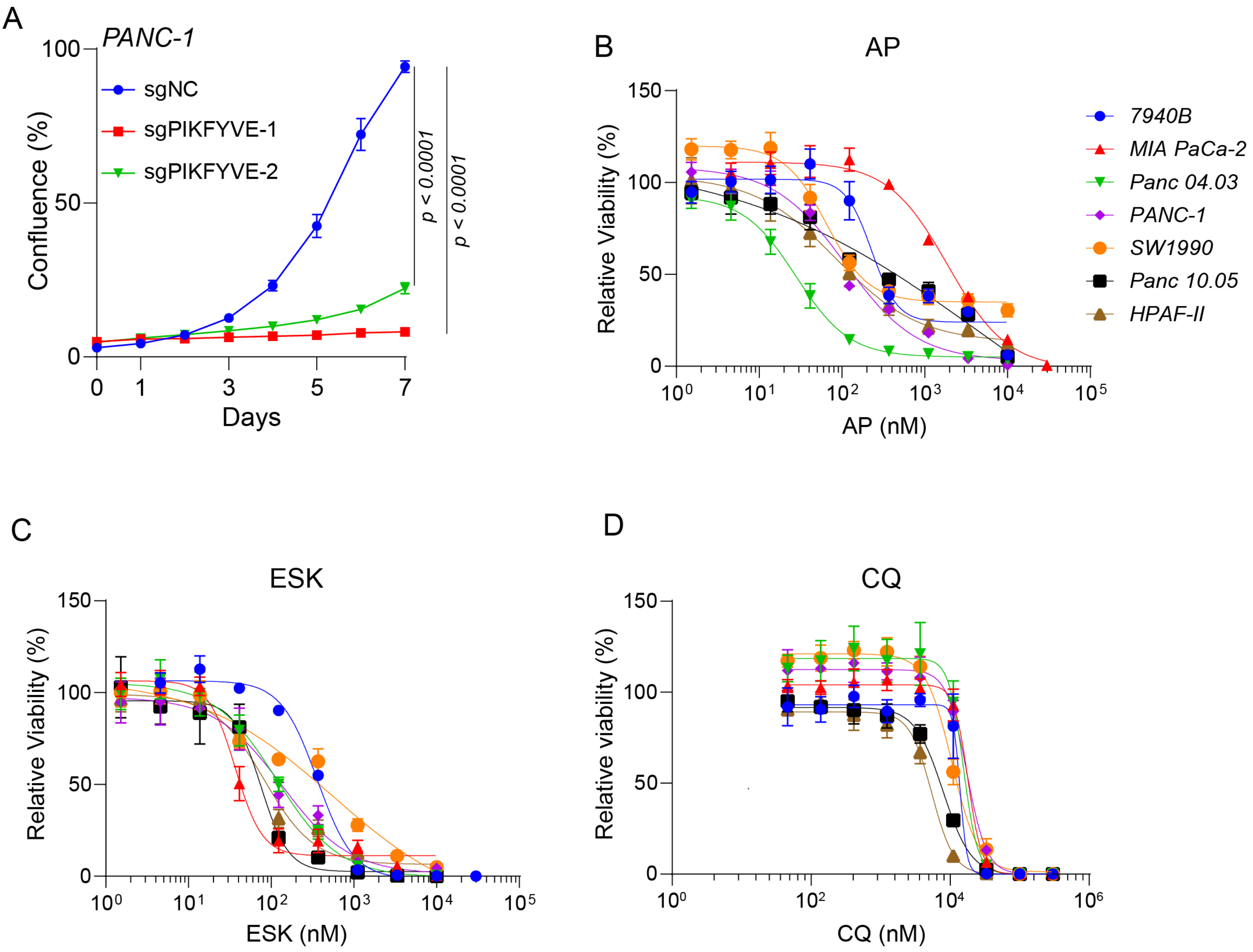


E

| | AP (nM) | ESK (nM) | CQ (nM) |
|---|---|---|---|
| *KPC-7940B* | 228.6 | 361.4 | 13893 |
| *MIA PaCa-2* | 1902 | 36.95 | 18019 |
| *Panc 04.03* | 27.97 | 117.8 | 15414 |
| *PANC-1* | 110.4 | 135.3 | 17306 |
| *SW 1990* | 65.19 | 554.7 | 10641 |
| *Panc 10.05* | 4960 | 75.22 | 7913 |
| *HPAF-II* | 85.82 | 81.96 | 5446 |

Extended Data Fig. 5

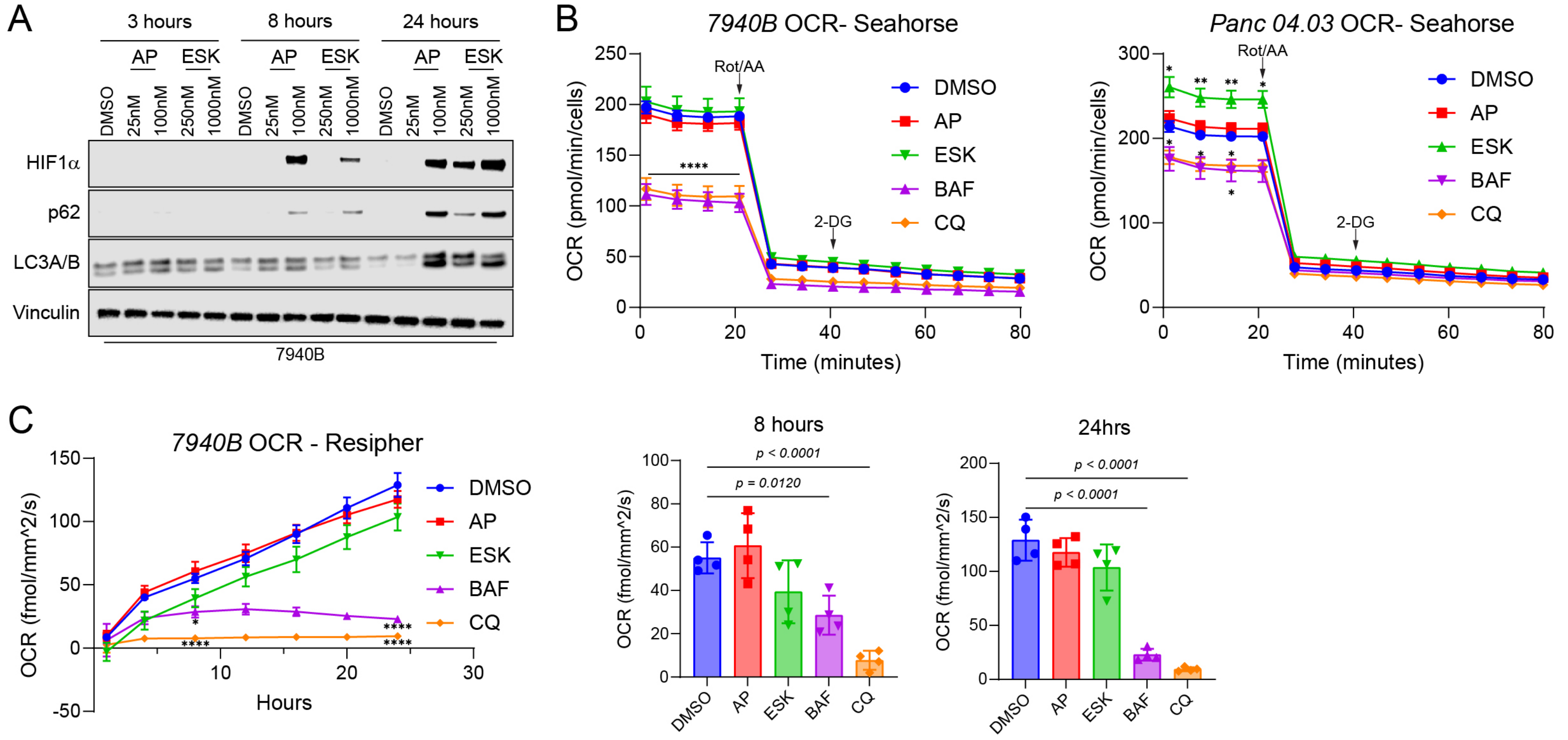


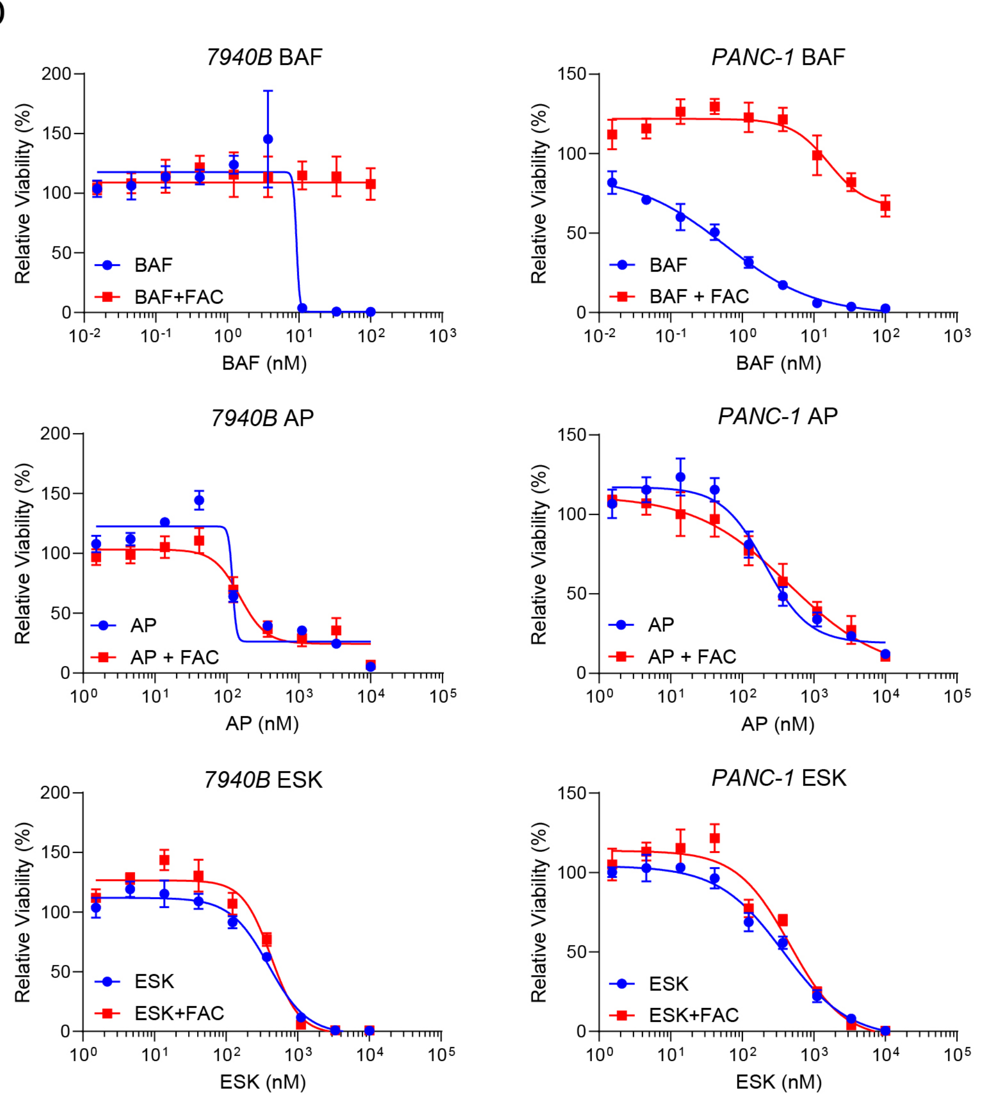


E

Half-maximal inhibitory concentration ($IC_{50}$)

| 7940B | -FAC | +FAC |
|---|---|---|
| BAF | 9.241 | N/A |
| AP | 120.1 | 151.8 |
| ESK | 395.7 | 425.9 |

| PANC-1 | -FAC | +FAC |
|---|---|---|
| BAF | 0.5775 | 16.3 |
| AP | 210.7 | 449.8 |
| ESK | 377.4 | 464.3 |

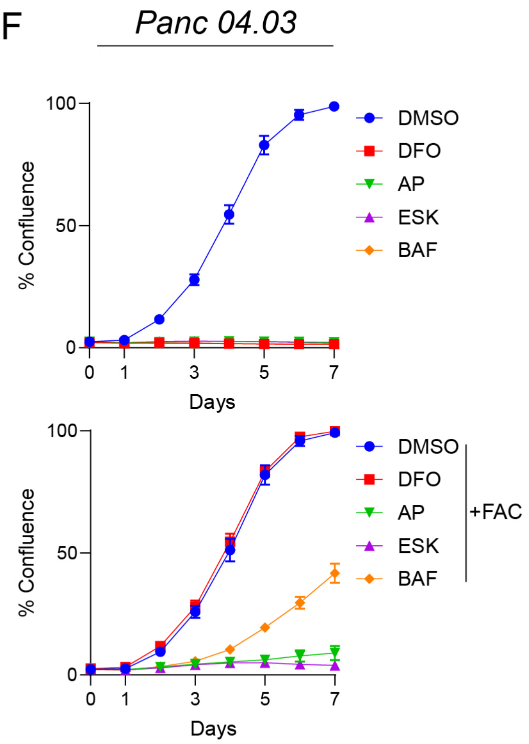

Extended Data Fig. 6

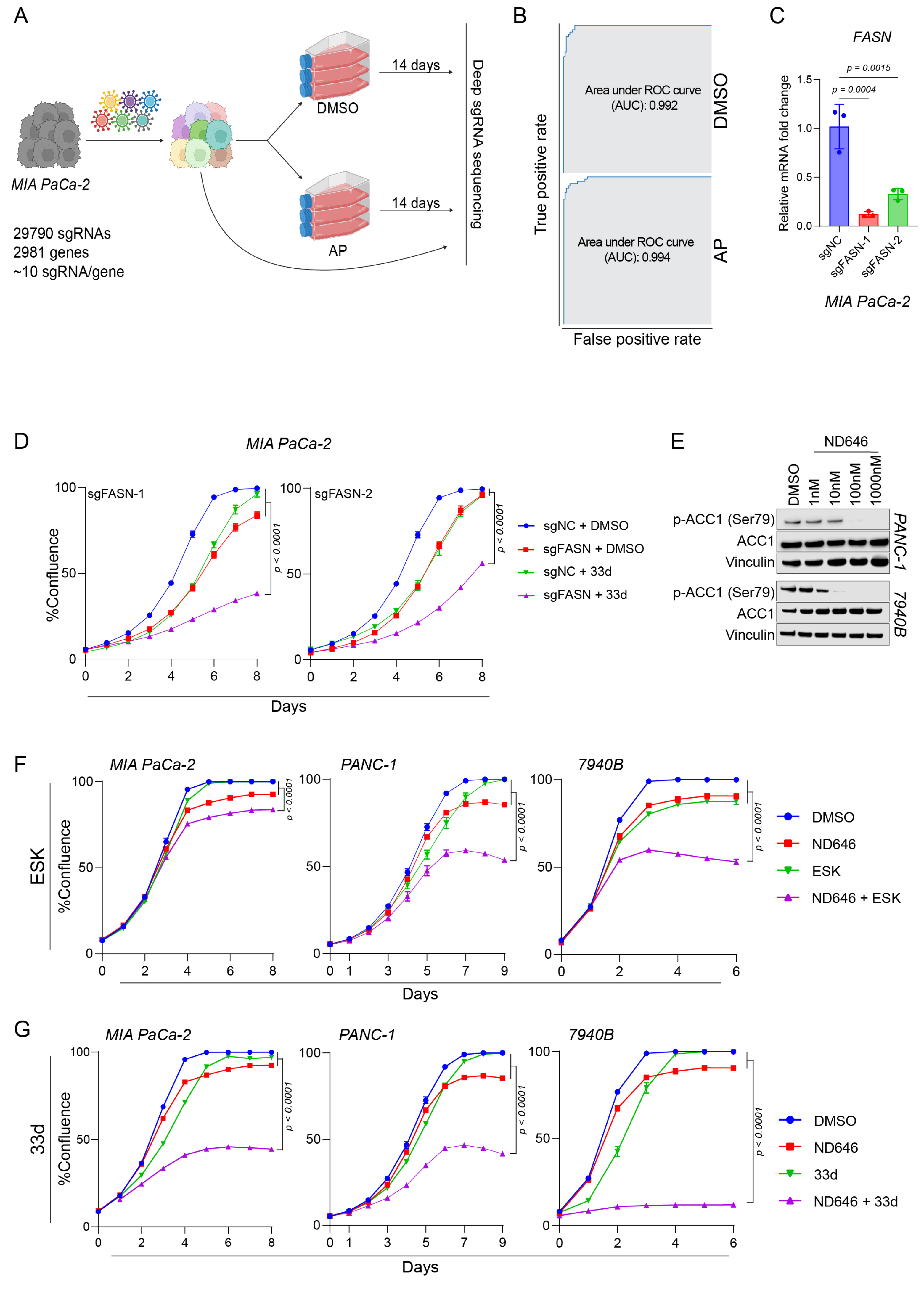

Extended Data Fig. 7

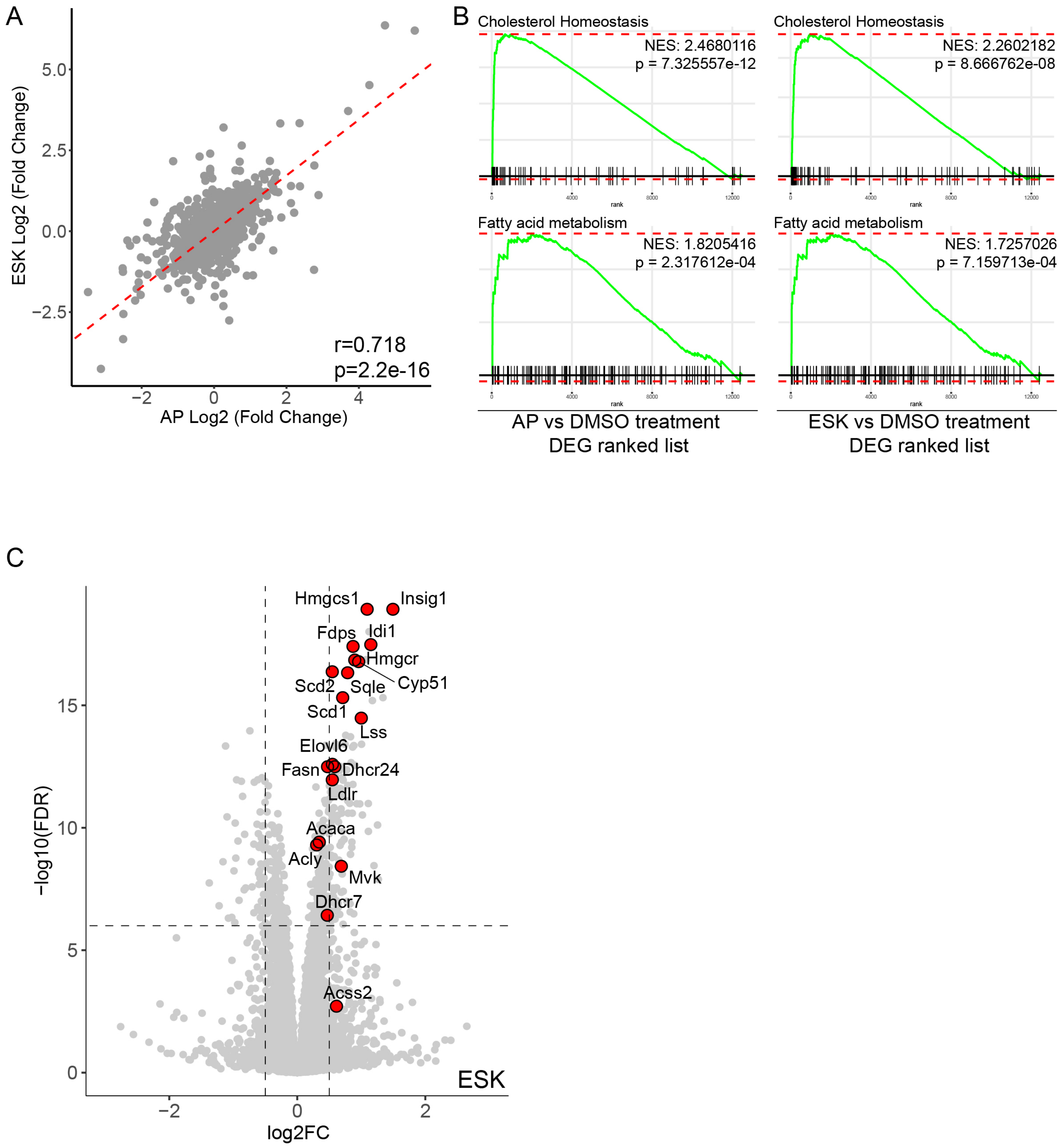

Extended Data Fig. 8

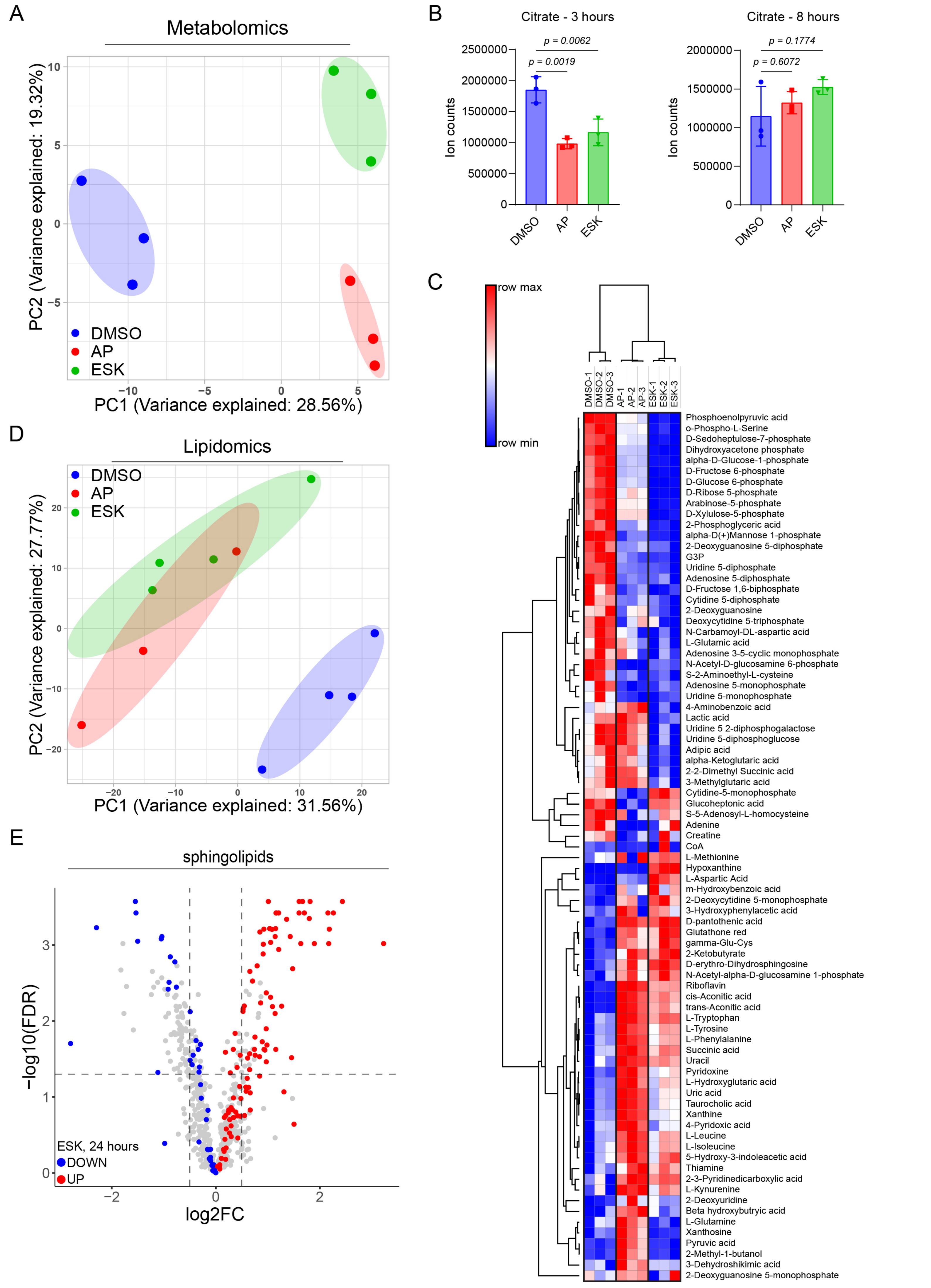

Extended Data Fig. 9

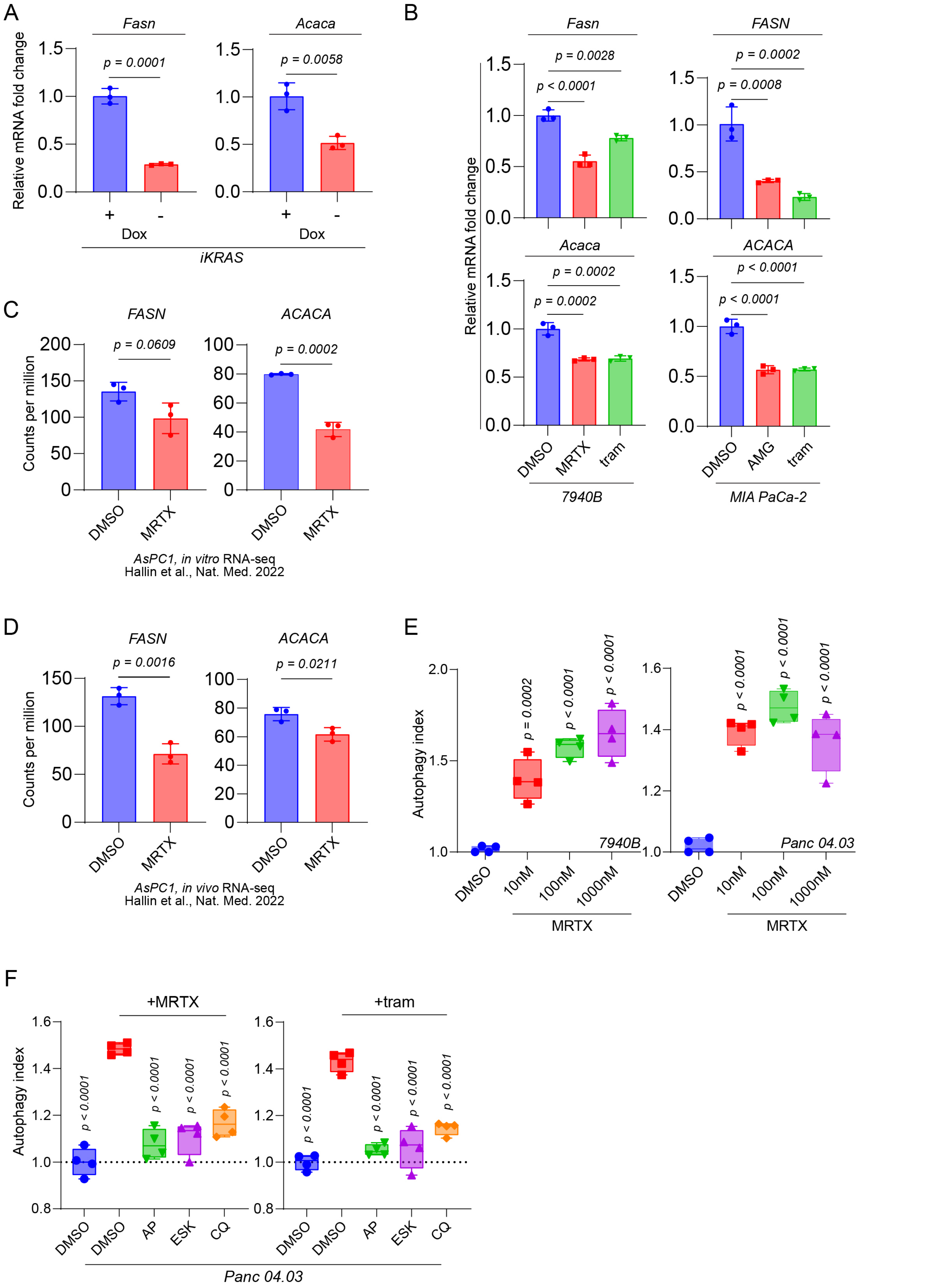

Extended Data Fig. 10

A

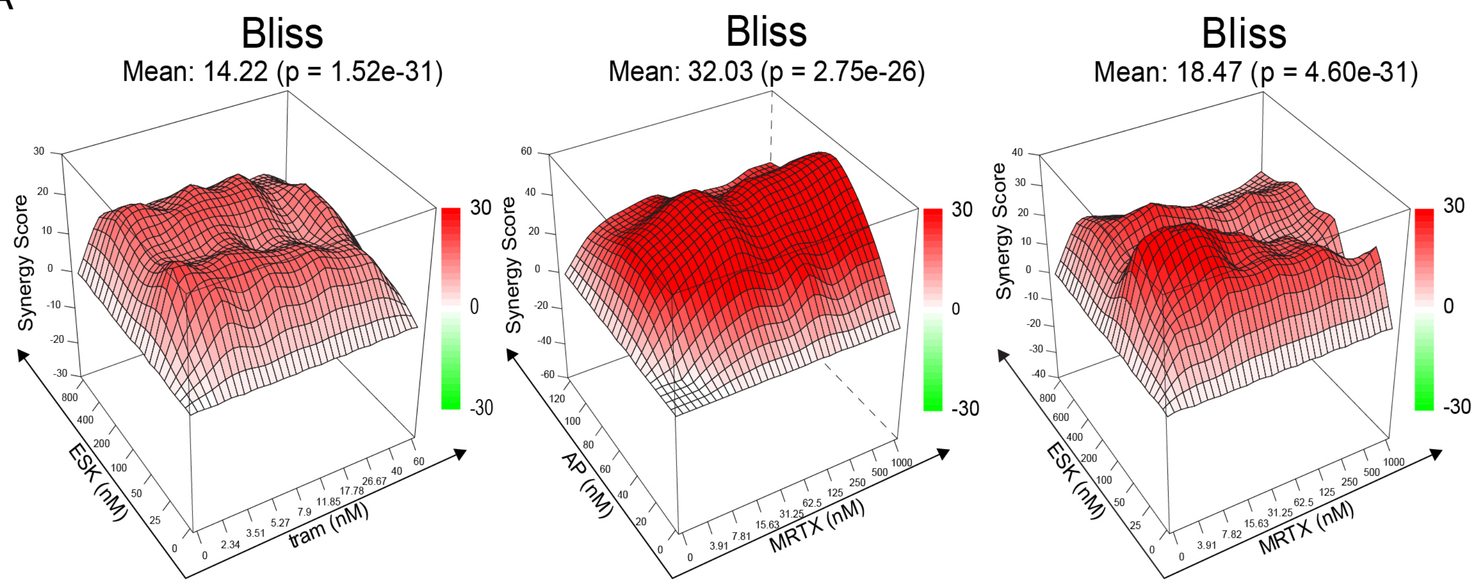


7940B

B

| tram (nm) \ ESK (nM) | 0 | 25 | 50 | 100 | 200 | 400 | 800 |
|---|---|---|---|---|---|---|---|
| 60 | 71 | 75 | 80 | 86 | 83 | 86 | 95 |
| 40 | 57 | 70 | 76 | 83 | 82 | 85 | 94 |
| 26.7 | 51 | 64 | 70 | 71 | 72 | 82 | 91 |
| 17.8 | 36 | 51 | 60 | 62 | 65 | 77 | 87 |
| 11.9 | 26 | 39 | 52 | 53 | 58 | 69 | 82 |
| 7.9 | 18 | 33 | 47 | 52 | 51 | 66 | 80 |
| 5.27 | 15 | 26 | 46 | 47 | 49 | 60 | 79 |
| 3.51 | 20 | 36 | 48 | 52 | 50 | 67 | 80 |
| 2.34 | 10 | 18 | 43 | 44 | 41 | 58 | 78 |
| 0 | 0 | -1 | 13 | 21 | 18 | 34 | 61 |

| MRTX (nM) \ AP (nM) | 0 | 20 | 40 | 60 | 80 | 100 | 120 |
|---|---|---|---|---|---|---|---|
| 1000 | 3 | 54 | 65 | 83 | 89 | 90 | 91 |
| 500 | 24 | 53 | 59 | 78 | 88 | 88 | 90 |
| 250 | 24 | 47 | 56 | 74 | 85 | 85 | 85 |
| 125 | 16 | 49 | 48 | 65 | 79 | 82 | 83 |
| 62.5 | 11 | 34 | 41 | 57 | 70 | 80 | 78 |
| 31.3 | -4 | 24 | 35 | 50 | 67 | 79 | 79 |
| 15.6 | -11 | 14 | 24 | 37 | 58 | 70 | 75 |
| 7.82 | -8 | 9 | 19 | 34 | 52 | 68 | 73 |
| 3.91 | -7 | -2 | 8 | 25 | 43 | 63 | 67 |
| 0 | 0 | 5 | -6 | 3 | 22 | 46 | 55 |

| MRTX (nM) \ ESK (nM) | 0 | 25 | 50 | 100 | 200 | 400 | 600 | 800 |
|---|---|---|---|---|---|---|---|---|
| 1000 | 28 | 44 | 41 | 53 | 74 | 81 | 89 | 91 |
| 500 | 32 | 44 | 55 | 49 | 72 | 82 | 90 | 89 |
| 250 | 27 | 45 | 58 | 52 | 68 | 82 | 87 | 87 |
| 125 | 23 | 38 | 53 | 48 | 63 | 77 | 85 | 85 |
| 62.5 | 15 | 32 | 50 | 47 | 61 | 77 | 83 | 83 |
| 31.3 | 5 | 20 | 39 | 42 | 56 | 74 | 80 | 80 |
| 15.6 | -6 | 10 | 39 | 37 | 55 | 72 | 82 | 81 |
| 7.82 | -9 | 0 | 34 | 36 | 52 | 67 | 82 | 82 |
| 3.91 | -12 | -4 | 21 | 29 | 49 | 59 | 80 | 82 |
| 0 | 0 | -8 | 12 | 9 | 40 | 47 | 65 | 70 |

7940B

C

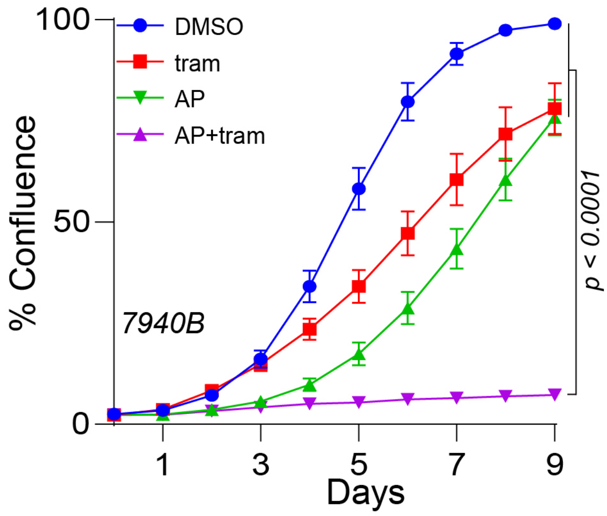

Extended Data Fig. 11

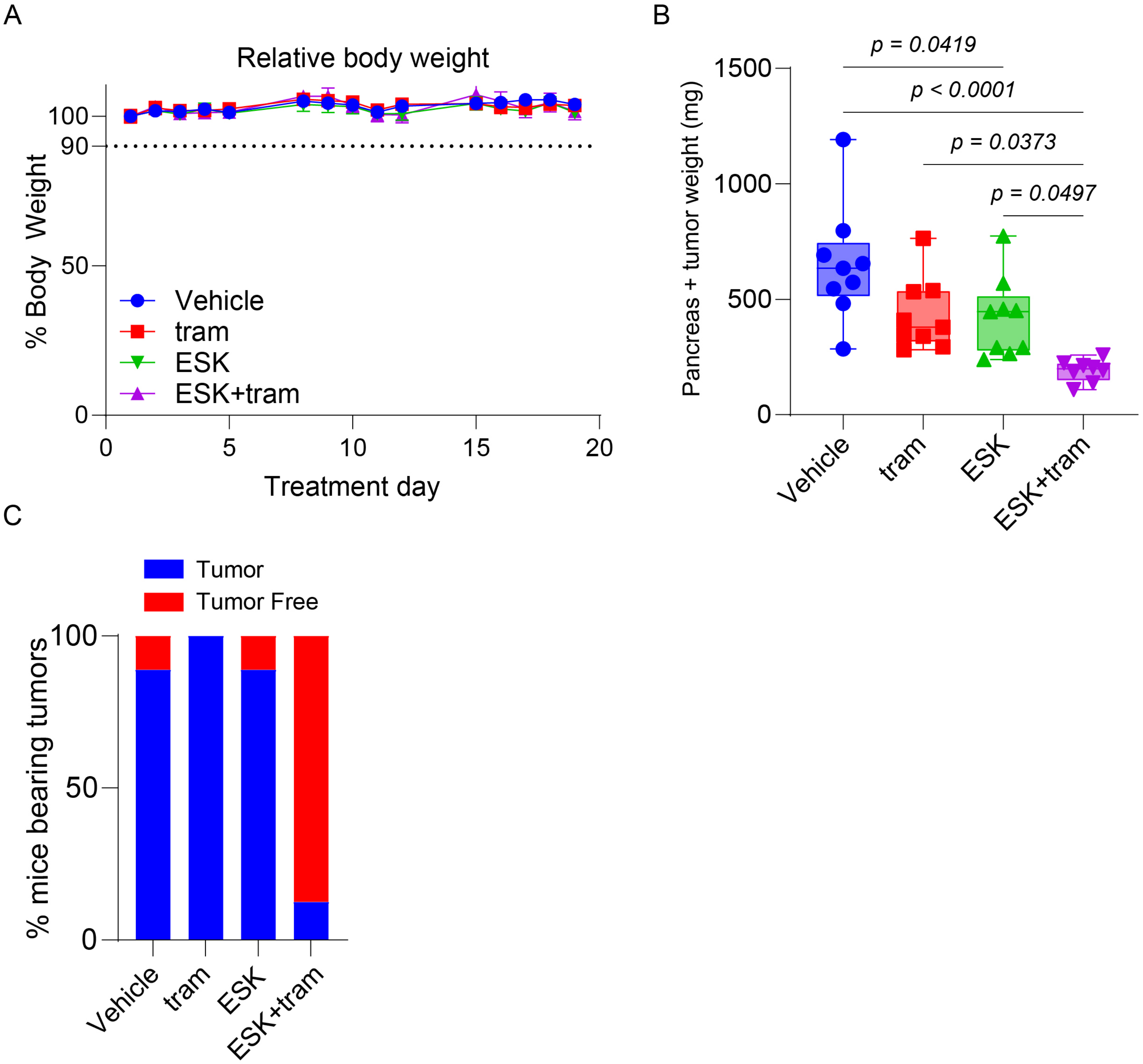
